# Knockout of Rab27b exacerbates neuropathology in alpha-synuclein mouse models

**DOI:** 10.64898/2025.12.19.695485

**Authors:** Kasandra Scholz, Mary A. Gannon, Lehmann Matheny, Rohma Syed, William J. Stone, Carley Craig, Roschongporn Ekkatine, Talene A. Yacoubian

## Abstract

Parkinson’s Disease (PD) and other synucleinopathies are characterized by the formation of inclusions comprised of alpha-synuclein (αsyn) among other proteins, but the mechanisms by which these inclusions form and cause toxicity are not well understood. We have previously reported that the small GTPase Rab27b modulates autophagic-lysosomal function in neurons and supports lysosomal degradation of αsyn across multiple αsyn cellular models. Knockout (KO) and knockdown of Rab27b damage lysosomal degradative capacity and exacerbate αsyn pathology, while Rab27b overexpression is conversely protective in cellular αsyn models. Elevations of Rab27b seen in human synucleinopathies suggest a compensatory role for Rab27b in these disorders. Here, we examined the role Rab27b plays *in vivo* in the context of both A53T genetic αsyn overexpression and viral AAV αsyn overexpression mouse models. Rab27b knockout in A53T^+^ mice did not alter motor behavior or survival. However, Rab27b knockout increased proteinase-K resistant αsyn in the cortex, striatum, and substantia nigra of A53T mice starting as early as six months of age. Additionally, Rab27b KO increased phosphorylated S129 αsyn in the cortex and nigra. Astrocyte and microglial activation were also observed upon Rab27b KO in the A53T model. In the AAV αsyn model, Rab27b KO resulted in accelerated dopaminergic cell loss in the nigra. Collectively, we report that loss of Rab27b results in elevated neuropathology in PD-relevant brain regions, validating its role as a therapeutic target in synucleinopathies.

## INTRODUCTION

Parkinson’s Disease (PD) is the most common movement disorder and second most common neurodegenerative disorder^1^. The hallmark neuropathological feature of PD and other synucleinopathies is the formation of Lewy bodies and neurites, which are enriched in alpha-synuclein (αsyn)^2^. αSyn aggregates within cells and is thought to be a key driver of disease pathology, although the biological mediators of its aggregation and toxicity are not fully characterized.

The Rab GTPase family is comprised of over 60 members, many of which are implicated in PD pathology^3–7^. Several of these proteins, including Rabs 1a, 3a, 8a, 11, and 39 have been linked specifically to αsyn aggregation and toxicity^8–11^. Proteins in this family canonically play roles in intracellular membrane trafficking, regulating a wide variety of processes that notably include autophagy and endolysosomal trafficking^12–16^. Endolysosomal and autophagic dysfunction are heavily implicated in PD and other proteinopathies, where they may contribute to disease pathogenesis and progression^17–21^. Both monomeric and oligomeric species of αsyn can be degraded by the lysosome^22–24^, and autophagic-lysosomal dysfunction can impair this process and facilitate unwanted protein aggregation^25–27^. Lysosomal dysfunction can exacerbate or contribute to αsyn pathology^28,29^, and several lysosomal genes including *GBA1* and *ATP13A2* are associated with elevated PD risk^30–33^. Proteins that modulate lysosomal function are therefore promising targets for understanding synucleinopathies and identifying new therapeutic strategies. We recently identified the Rab27b GTPase as a modulator of lysosomal function and αsyn aggregation across multiple models^34,35^. Rab27b knockout (KO) in mice impairs lysosomal axonal trafficking in primary neurons and lysosomal enzymatic maturation and activity in both primary neurons and brain lysates^35^. Rab27b KO additionally exacerbates the accumulation of pS129-phosphorylated αsyn in primary neurons treated with αsyn pre-formed fibrils^35^. Similarly, knockdown (KD) of Rab27b in a human doxycycline-inducible αsyn cell line damages lysosomal function, increases αsyn accumulation in lysosomes, and exacerbates αsyn pathology, while Rab27b overexpression (OE) is protective in this model^34,35^. Protein levels of Rab27b are increased in human brain lysates from patients with PD and Dementia with Lewy Bodies (DLB), and increased Rab27b levels are also observed in Incidental Lewy Body Disease (iLBD), which is viewed as potentially the earliest stages of synucleinopathy^34,35^. These elevations in human disease may reflect a compensatory response in which the brain is attempting to boost lysosomal degradative function in the setting of proteostatic stress due to αsyn aggregation.

While these studies show that loss of Rab27b increases αsyn pathology in two *in vitro* αsyn models, here we examine whether Rab27b KO impacts αsyn pathology and neuronal loss in two *in vivo* αsyn models: the A53T human αsyn-overexpressing transgenic mouse model^36^ and the adeno-associated virus (AAV) αsyn overexpression model^37^. We found that Rab27b KO in the A53T mice exacerbated neuropathology, including insoluble αsyn accumulation and elevated astrocytic and microglial gliosis. We also found accelerated dopaminergic neuron loss in the AAV αsyn model. Our findings support a role for Rab27b in αsyn pathology *in vivo*, highlighting it as a potential therapeutic target in synucleinopathies.

## RESULTS

### Rab27b KO does not exacerbate wire hang or survival defects in the A53T^+^ model

In humans, the A53T αsyn mutation is a rare autosomal dominant mutation associated with early disease onset and increased αsyn aggregation^38,39^. The A53T^+^ transgenic mouse strain overexpresses A53T mutant αsyn and exhibits insoluble αsyn accumulation, astrogliosis, and severe motor impairments that rapidly progress to death^36,40^. To test the hypothesis that Rab27b KO could exacerbate the characteristic behavioral and pathologic changes seen in A53T^+^ αsyn mice, we crossbred A53T^+^ transgenic and Rab27b KO mouse lines to generate four final genotypes: WT (Rab27^+/+^ A53T^-/-^), A53T^+^ (Rab27b^+/+^ A53T^+/-^), Rab27b KO (Rab27b^-/-^ A53T^-/-^), and Rab27b KO/A53T^+^ (Rab27b^-/-^ A53T^+/-^) (Fig. 1).

**Figure 1:**
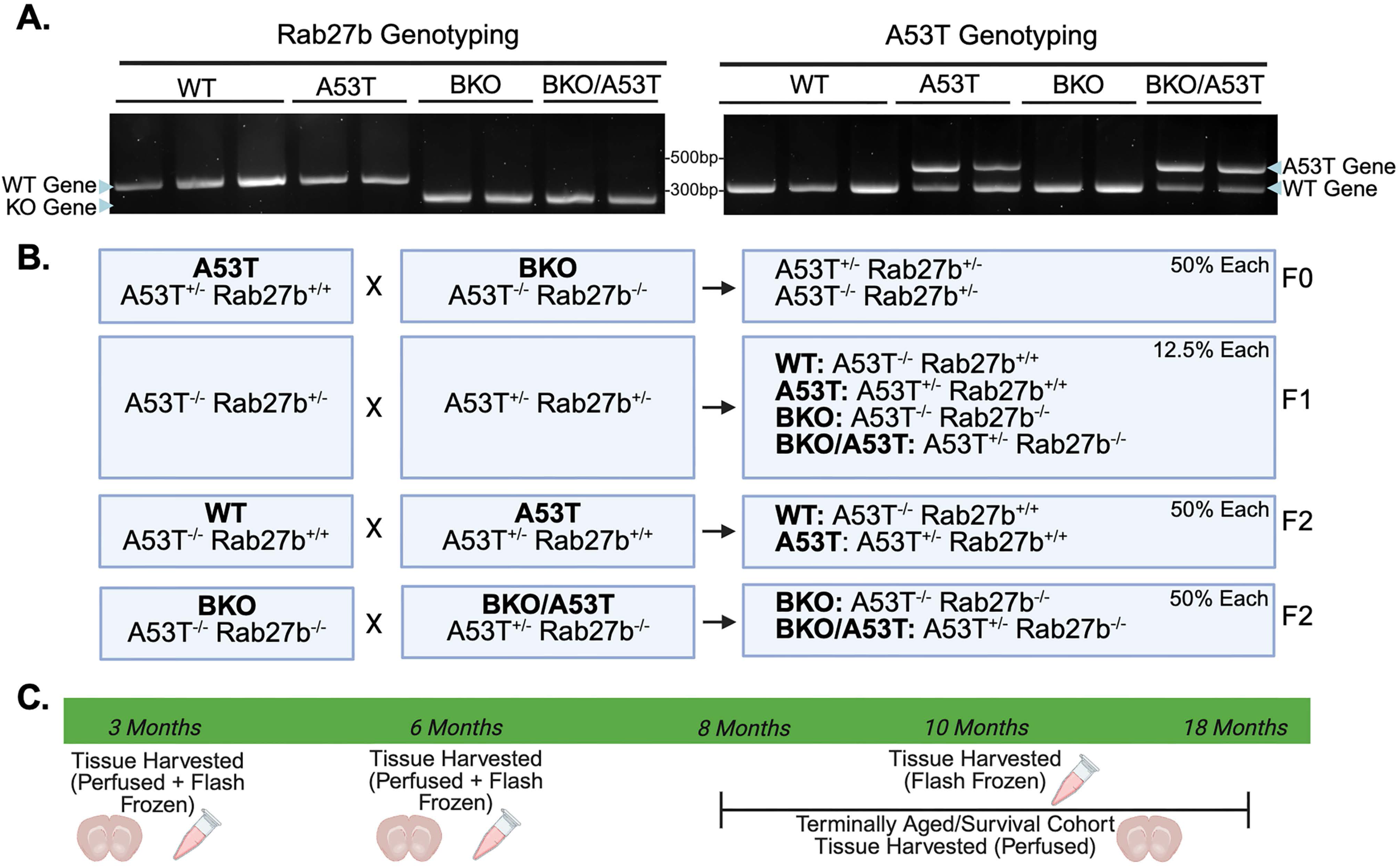
Mouse model validation, breeding schematic, and cohort timeline. a: Genotyping validates Rab27b knockout (left) and the presence of the A53T mutation (right) in the Rab27b KO/A53T^+^ model. b: Summary of breeding strategy to generate experimental mice used in this study, including WT, Rab27b KO, A53T^+^, and Rab27b KO/A53T^+^ mice. c: Timeline summarizing ages of animals at time of tissue collection.

We first assessed motor behavior. A53T^+^ transgenic mice develop severe motor impairments as they age, exhibiting progressive paralysis that eventually leads to death^36^. We assessed two separate cohorts of animals (a younger 3-4 month cohort and an older 7-10 month cohort) using a four paw wire hang test^41^. While other strains of mice expressing human A53T mutant αsyn exhibit motor defects as early as one month of age^42,43^, the present A53T^+^ strain does not typically show early motor defects. For example, one group found G2-3 A53T^+^ mice show no defects in rotor rod or pole test performance at six months of age^44^. In contrast to these previously reported findings, we detected motor defects in A53T^+^ mice on the wire hang test beginning at four months of age (Fig. 2b).

**Figure 2:**
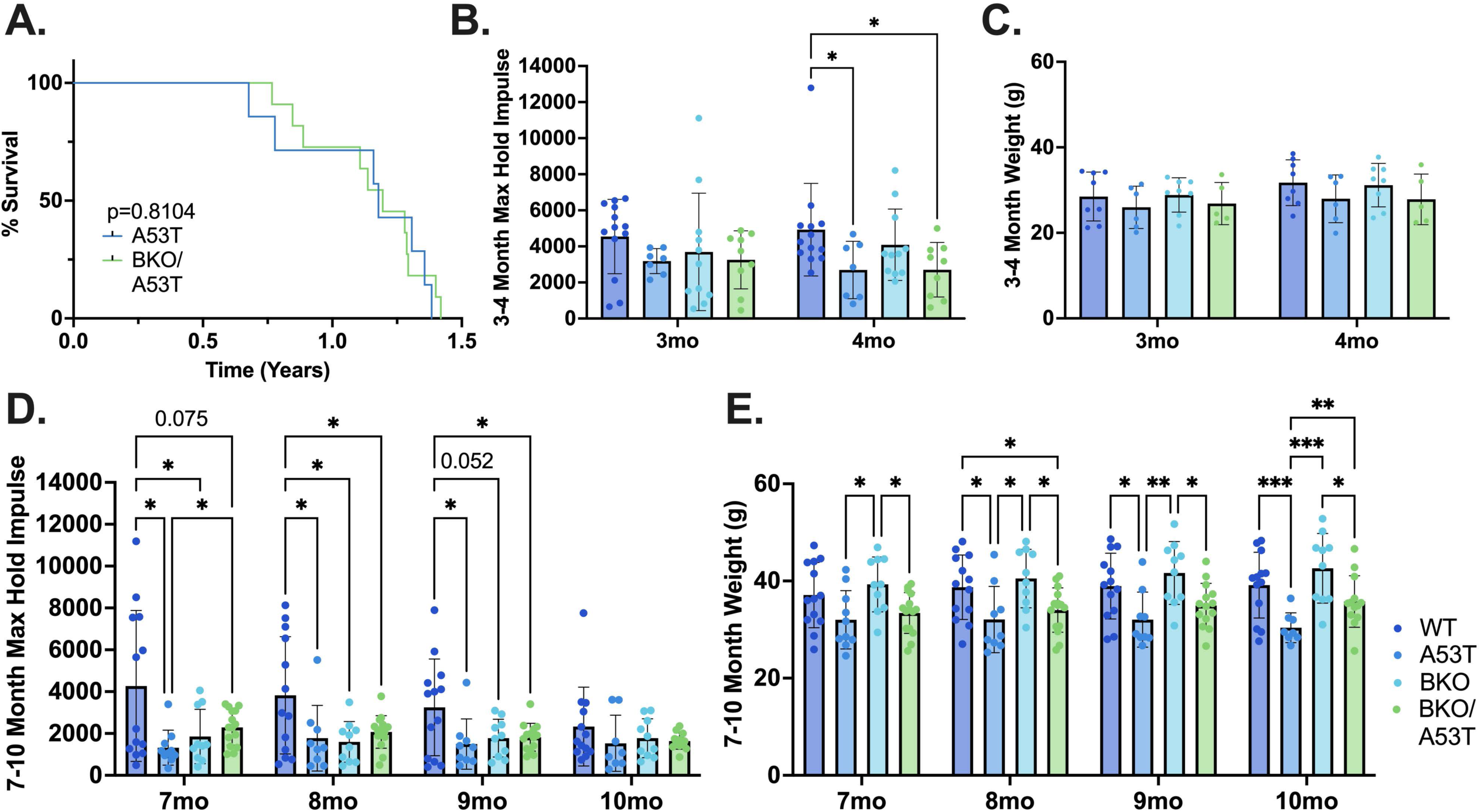
Rab27b KO does not exacerbate survival or wire hang capacity in A53T^+^ mice. **a:** Survival curve showing age of onset of terminal spinal paralysis in A53T^+^ animals. Log-rank (Mantel-Cox) test. N=7-11 individual mice per group. b-c: Maximum holding impulse (b) and weights (c) of young cohort animals at three- and four-months of age. Two-way repeated measures ANOVA followed by Fisher’s LSD post-hoc test; *p<0.05. N=7-13 individual mice per group. d-e: Maximum holding impulse (d) and weights (e) of aged cohort animals at seven to 10 months. Mixed effects model (REML) followed by Fisher’s LSD post-hoc test; *p<0.05, **p<0.01, ***p<0.001. N=8-15 individual mice per group. All error bars denote standard deviation from the mean.

In the younger behavioral mouse cohort, both A53T^+^ and Rab27b KO/A53T^+^ groups showed reduced wire hang capacity relative to WT animals at four months of age, but there were no differences between the A53T^+^ mice and Rab27b KO/A53T^+^ mice (Fig. 2b). No differences in weight were noted between the different genotypes in the young cohort (Fig. 2c). Similarly, in the older behavioral cohort we found that both A53T^+^ and Rab27b KO/A53T^+^ mice had significantly reduced wire hang performance relative to WT mice at eight and nine months of age (Fig. 2d). A53T^+^ mice showed reduced weights as they aged (Fig. 2e), and we adjusted for weight differences in the maximum holding impulse calculations. By 10 months, there were no differences between any groups, likely due to a decline in the performance of the WT animals (Fig. 2d; WT 7mo vs 10mo p=0.016). Surprisingly, the Rab27b KO/A53T^+^ mice performed better than the A53T^+^ mice at seven months, but this was not noted in any other time points or in the younger cohort (Fig. 2b,d). At all other time points, there were no significant differences between the A53T^+^ and Rab27b KO/A53T^+^ mice. Interestingly, we found that the Rab27b KO mice also demonstrated wire hang defects at seven and eight months of age, indicating that Rab27b KO alone can impact motor behavior (Fig. 2d).

We also assessed longevity in A53T^+^ compared to Rab27b KO/A53T^+^ mice to determine if Rab27b KO would impact survival. Animals were examined daily and sacrificed upon onset of severe hind-limb paralysis. Approximately 10% of A53T^+^ transgenic mice are non-penetrant and fail to exhibit motor deficits even at advanced ages^36^, and animals that reached 18 months of age without exhibiting motor deficits were excluded from the survival analysis. None of the WT or Rab27b KO mice exhibited paralysis, and we found no differences in survival between A53T^+^ and Rab27b KO/A53T^+^ mice (Fig. 2a).

### Rab27b KO/A53T^+^ mice show autophagic-lysosomal dysregulation

We have previously found that while Rab27b KO impairs lysosomal enzymatic activity and increases pre-formed αsyn fibril-mediated pathology in primary neurons, it does not result in measurable changes in the autophagic substrate marker p62 or the lysosomal marker LAMP1 in the absence of αsyn overexpression^35^. Rab27b KO/A53T^+^ mice showed significantly increased p62 protein levels in cortical lysates compared to WT mice at six months of age and compared to all other groups at 10 months of age (Fig. 3a-b). Similarly, Rab27b KO/A53T^+^ mice also had increased LAMP1 levels at 10 months in cortical lysates compared to all other groups (Fig. Xa-b). Younger, three-month-old Rab27b KO/A53T^+^ animals did not exhibit changes in these markers compared to any other group (Fig. 3a-b). These findings provide evidence that Rab27b loss contributes to autophagic-lysosomal dysregulation with age in this αsyn model.

**Figure 3:**
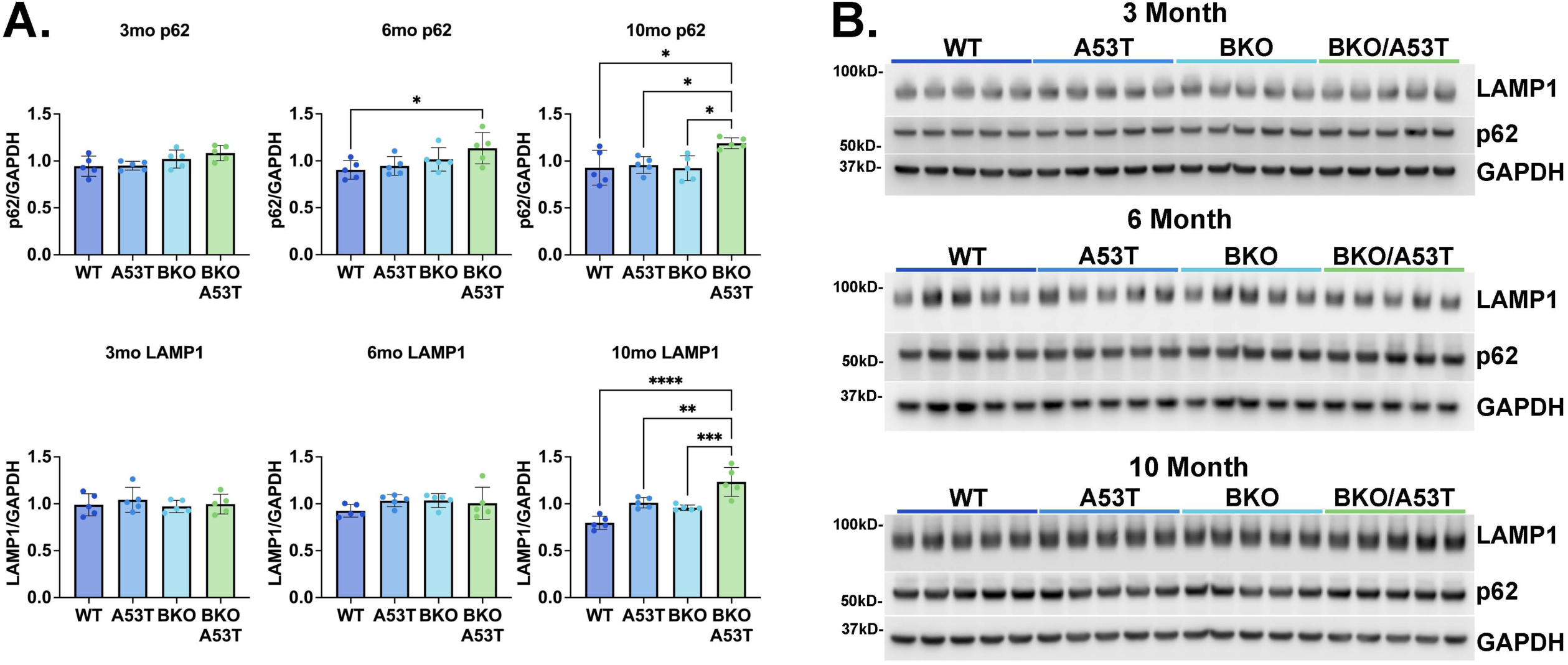
Rab27b KO induces dysregulation in autophagic-lysosomal markers in A53T^+^ mice. **a:** Quantification of p62 (top) and LAMP1 (bottom) levels in the cortex of WT, A53T^+^, Rab27b KO, and Rab27b KO/A53T^+^ mice at three-, six-, and 10-months of age. One-way ANOVA followed by Tukey’s multiple comparisons post-hoc test; *p<0.05, **p<0.01, ***p<0.001, ****p<0.0001. N=5 individual mice per group. b: Representative Western blots for p62, LAMP1, and GAPDH in the cortex of WT, A53T^+^, Rab27b KO, and Rab27b KO/A53T^+^ mice at three-, six-, and 10-months of age. All error bars denote standard deviation from the mean. Note that the six and 10 month GAPDH bands overlap between this figure and Fig. 8, as markers were probed on the same membranes.

**Figure 4:**
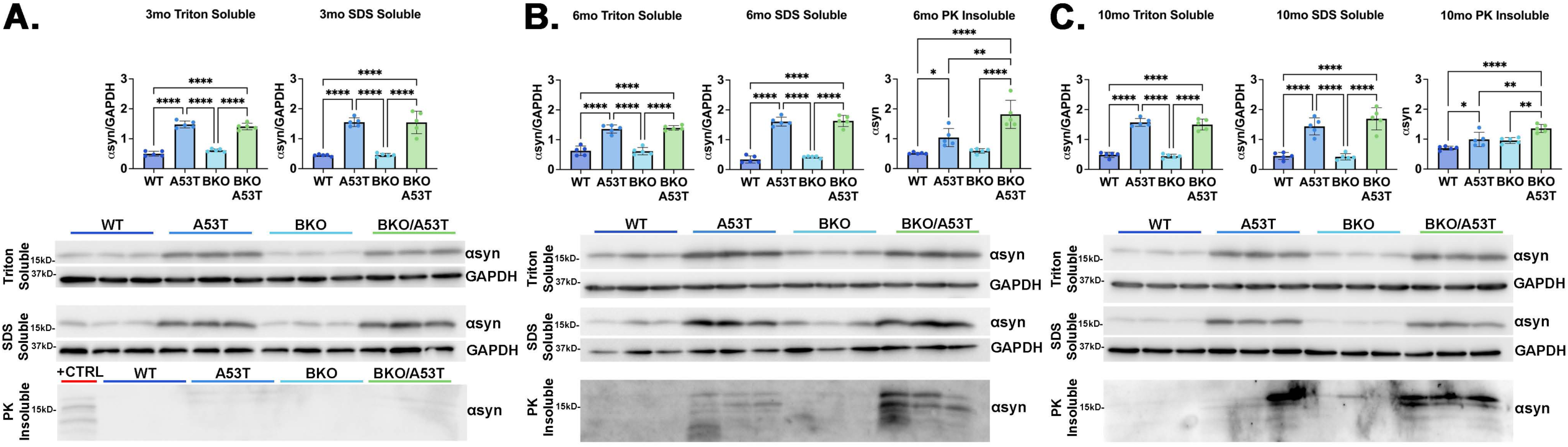
Rab27b KO results in elevated accumulation of PK-insoluble **α**syn in cortical lysates. a: Representative Western blots and quantification of Triton- and SDS-soluble αsyn levels in the cortex of three-month-old WT, A53T^+^, Rab27b KO, and Rab27b KO/A53T^+^ mice. One-way ANOVA followed by Tukey’s multiple comparisons post-hoc test; ****p<0.0001. Positive control in PK-digested blot is cortical lysate from a 10-month-old Rab27b KO/A53T^+^ animal. N=5 individual mice per group. b: Representative Western blots and quantification of Triton- and SDS-soluble and PK-insoluble αsyn levels in the cortex of six-month-old WT, A53T^+^, Rab27b KO, and Rab27b KO/A53T^+^ mice. One-way ANOVA followed by Tukey’s multiple comparisons post-hoc test; *p<0.05, **p<0.01, ****p<0.0001. N=5 individual mice per group. c: Representative Western blots and quantification of Triton- and SDS-soluble and PK-insoluble αsyn levels in the cortex of 10-month-old WT, A53T^+^, Rab27b KO, and Rab27b KO/A53T^+^ mice. One-way ANOVA followed by Tukey’s multiple comparisons post-hoc test; *p<0.05, **p<0.01, ****p<0.0001. N=5 individual mice per group. All error bars denote standard deviation from the mean.

### Rab27b KO exacerbates the accumulation of proteinase K-insoluble alpha-synuclein

We next assessed the accumulation of insoluble αsyn in cortical lysates from three-, six-, and 10-month-old animals. Insoluble αsyn accumulation is a hallmark of pathology in A53T^+^ mice^36^, and we wanted to determine if Rab27b KO and concurrent lysosomal deficits could increase pathologic αsyn in the A53T^+^ model. Cortical lysates were serially fractionated using Triton X-100, SDS, and proteinase K (PK) digestion to isolate increasingly insoluble αsyn species and then assessed via Western blot. As anticipated, we found that the A53T^+^ and Rab27b KO/A53T^+^ groups had elevated αsyn levels relative to the WT and Rab27b KO animals in Triton X-100 and SDS soluble fractions at all time points (Fig. 2a-c). We were unable to detect quantifiable levels of PK-insoluble αsyn in the three-month-old cohort (Fig. 2a). While we did not see differences in αsyn levels between the A53T^+^ and Rab27bKO/A53T^+^ animals in the Triton X-100 or SDS soluble fractions, Rab27b KO/A53T^+^ mice showed significantly elevated levels of PK-resistant αsyn in the cortex at both six and 10 month time points compared to WT, A53T^+^, and Rab27b KO mice (Fig. 2b-c).

To validate this finding, we assessed total and PK-resistant αsyn levels in perfused tissue via immunohistochemistry (IHC) after PK digestion in separate three-month-old, six-month-old, and terminally aged cohorts (see Table 2 for terminally aged cohort demographics). In all cohorts, αsyn intensity was assessed in the sensorimotor cortex (Ctx) and dorsolateral striatum (Str). At all three time points, A53T^+^ and Rab27b KO/A53T^+^ animals again showed elevated total αsyn intensity relative to WT and Rab27b KO mice in non-PK-digested Ctx and Str (Fig. S1; 5a-f). Three-month-old animals did not display differences in PK-resistant αsyn in the Ctx or Str between any groups (Fig. S1). In the six-month-old mice, Rab27b KO/A53T^+^ animals displayed elevated PK-insoluble αsyn compared to all other groups in both the Ctx and Str (Fig. 5a-b). In the terminally aged cohort, Rab27bKO/A53T^+^ animals displayed elevated PK-insoluble αsyn compared to WT and Rab27b KO mice in both the Ctx and Str (Fig. 5c-d). Similarly, Rab27bKO/A53T^+^ mice additionally showed elevated PK-insoluble αsyn in the substantia nigra pars compacta (SNc) relative to the WT and Rab27b KO groups in the terminally aged cohort (Fig. 5e-f).

**Figure 5:**
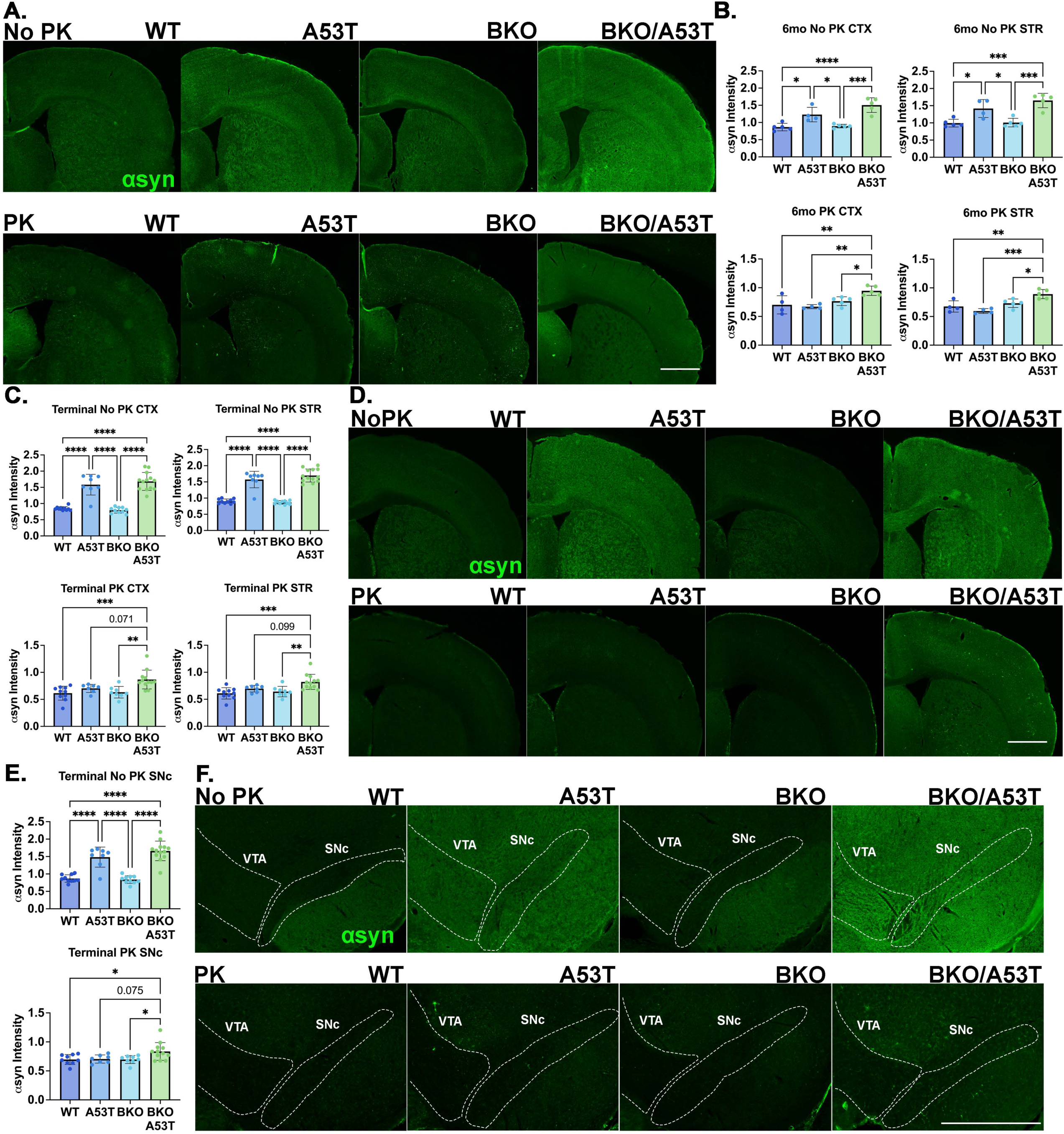
Rab27b KO increases accumulation of PK-insoluble **α**syn in the Ctx, Str, and SNc. a-b: Representative images (a) and quantification (b) of αsyn immunoreactivity in the Ctx and Str from brain tissue with and without PK digestion from six-month-old WT, A53T^+^, Rab27b KO, and Rab27b KO/A53T^+^ mice. One-way ANOVA followed by Tukey’s multiple comparisons post-hoc test; *p<0.05, **p<0.01, ***p<0.001, ****p<0.0001. N=4-5 individual mice per group. c-d: Quantification (c) and representative images (d) of αsyn immunoreactivity in the Ctx and Str from brain tissue with and without PK digestion from terminally aged WT, A53T^+^, Rab27b KO, and Rab27b KO/A53T^+^ mice. One-way ANOVA followed by Tukey’s multiple comparisons post-hoc test; **p<0.01, ***p<0.001, ****p<0.0001. N=8-13 individual mice per group. e-f: Quantification (e) and representative images (f) of αsyn immunoreactivity in the SNc from brain tissue with and without PK digestion from terminally aged WT, A53T^+^, Rab27b KO, and Rab27b KO/A53T^+^ mice. VTA and SNc were outlined using TH staining (not pictured). One-way ANOVA followed by Tukey’s multiple comparisons post-hoc test; *p<0.05, ****p<0.0001. N=8-13 individual mice per group. Ctx = Sensorimotor Cortex; Str = Dorsolateral Striatum; SNc = Substantia Nigra pars Compacta; VTA = Ventral Tegmental Area. Scale bars are 1000µm. All error bars denote standard deviation from the mean.

We finally assessed the accumulation of pathogenic αsyn marked by S129 phosphorylation (pS129) in PK-digested tissue in our terminally aged cohort. WT or Rab27b KO mice did not demonstrate any pS129-positive inclusions, but Rab27b KO/A53T^+^ mice showed significant elevations in pS129 αsyn signal relative to A53T^+^ animals in both the Ctx and SNc (Fig. 6a-c). Collectively, we report that knockout of Rab27b can exacerbate the accumulation of pathologic αsyn across several brain regions.

**Figure 6:**
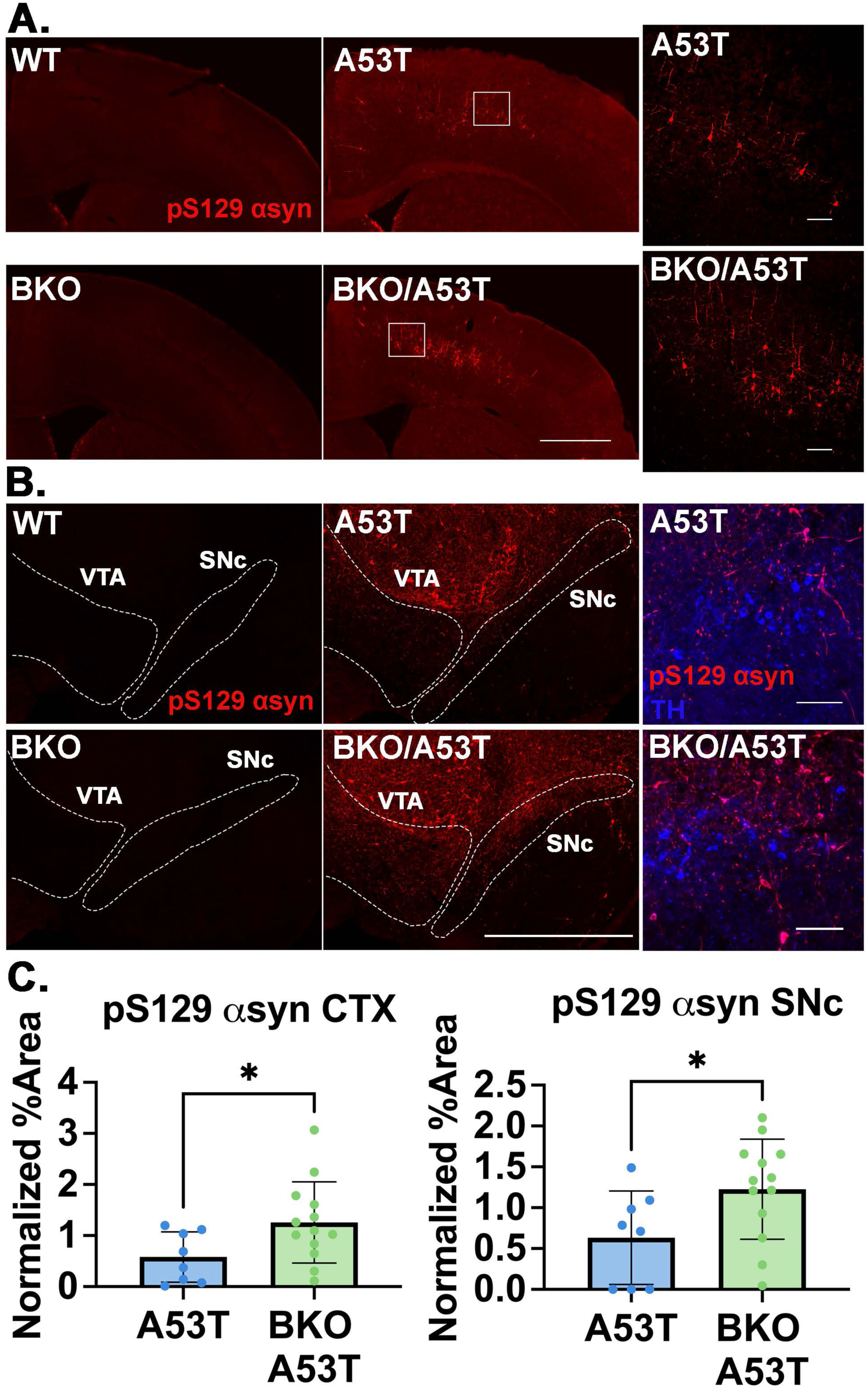
Rab27b KO exacerbates the accumulation of pS129. α**syn in A53T^+^ mice.** a: Representative images showing pS129 αsyn signal in the Ctx of terminally aged WT, Rab27b KO, A53T^+^, and Rab27b KO/A53T^+^ mice. Box shows area zoomed in on the right. b: Representative images showing pS129 αsyn signal in the SNc of terminally aged WT, Rab27b KO, A53T^+^, and Rab27b KO/A53T^+^ mice. VTA and SNc were outlined using TH staining (not pictured). Right images show zoomed in portion within the SNc along with TH staining. c: Quantification of area positive for pS129 αsyn in the Ctx and SNc. Student’s t-test; *p<0.05. N=8-13 individual mice per group. Ctx = Sensorimotor Cortex; SNc = Substantia Nigra pars Compacta; VTA = Ventral Tegmental Area. Scale bars on full-sized images are 1000µm. Scale bars on enlarged images are 100µm. All error bars denote standard deviation from the mean.

### Rab27b KO exacerbates neuroinflammation

Astrocytic gliosis is another major pathological characteristic seen in A53T^+^ αsyn mice, although it is typically restricted to the hindbrain^36^. To determine if Rab27b KO could elicit astrogliosis in additional brain regions, we stained three-month-old, six-month-old, and terminally aged mouse brains for the reactive astrocyte marker GFAP. At six months, we found significantly elevated GFAP signal in Rab27b KO/A53T^+^ mice in the Ctx compared to WT and Rab27b KO mice and in the Str compared to all other groups (Fig. S2a). At terminal ages, Rab27b KO/A53T^+^ showed increased GFAP signal in the Ctx, Str, and SNc compared to all other groups (Fig. 7a-b). To validate this finding, we measured GFAP levels in total protein lysates from the cortex at three, six, and 10 months of age by Western blotting. GFAP levels were elevated in Rab27b KO/A53T^+^ mice compared to all other groups at both six and 10 months but not in younger three-month-old animals (Fig. 8a-b).

**Figure 7:**
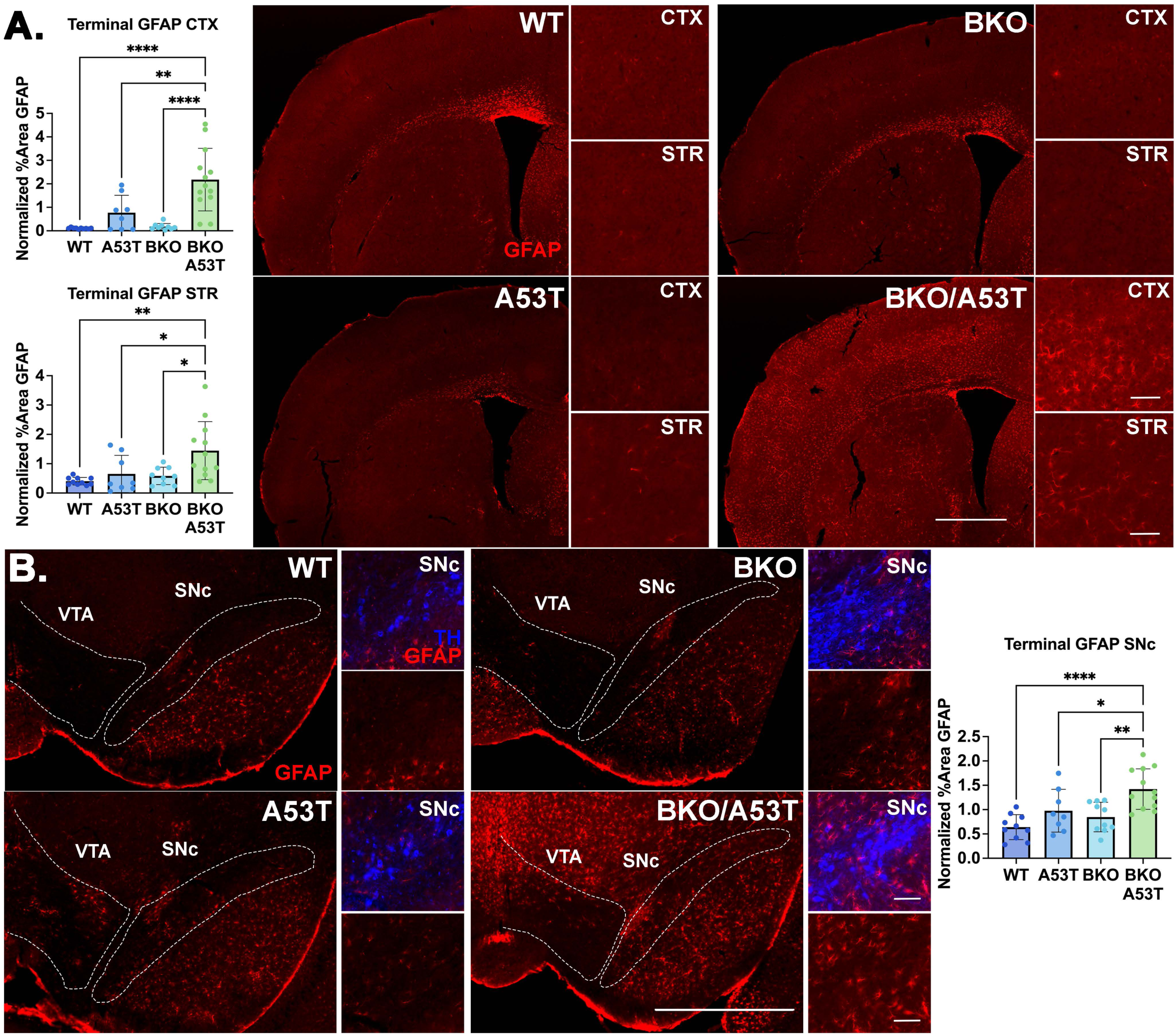
Rab27b KO/A53T^+^ mice show elevated GFAP signal in tissue from terminally aged mice. **a:** Quantification and representative images showing GFAP immunoreactivity in the Ctx and Str of terminally aged WT, Rab27b KO, A53T^+^, and Rab27b KO/A53T^+^ mice. Small panels are zoomed images. One-way ANOVA followed by Tukey’s multiple comparisons post-hoc test; *p<0.05, **p<0.01, ****p<0.0001. N=8-13 individual mice per group. b: Quantification and representative images showing GFAP immunoreactivity in the SNc of terminally aged WT, Rab27b KO, A53T^+^, and Rab27b KO/A53T^+^ mice. VTA and SNc were outlined using TH staining (not pictured). One-way ANOVA followed by Tukey’s multiple comparisons post-hoc test; *p<0.05, **p<0.01, ****p<0.0001. N=8-12 individual mice per group. Ctx = Sensorimotor Cortex; Str = Dorsolateral Striatum; SNc = Substantia Nigra pars Compacta; VTA = Ventral Tegmental Area. Scale bars on full-sized images are 1000µm. Scale bars on enlarged images are 100µm. All error bars denote standard deviation from the mean. Note that GFAP and Iba1 were stained on the same sections (see Fig. 9).

**Figure 8:**
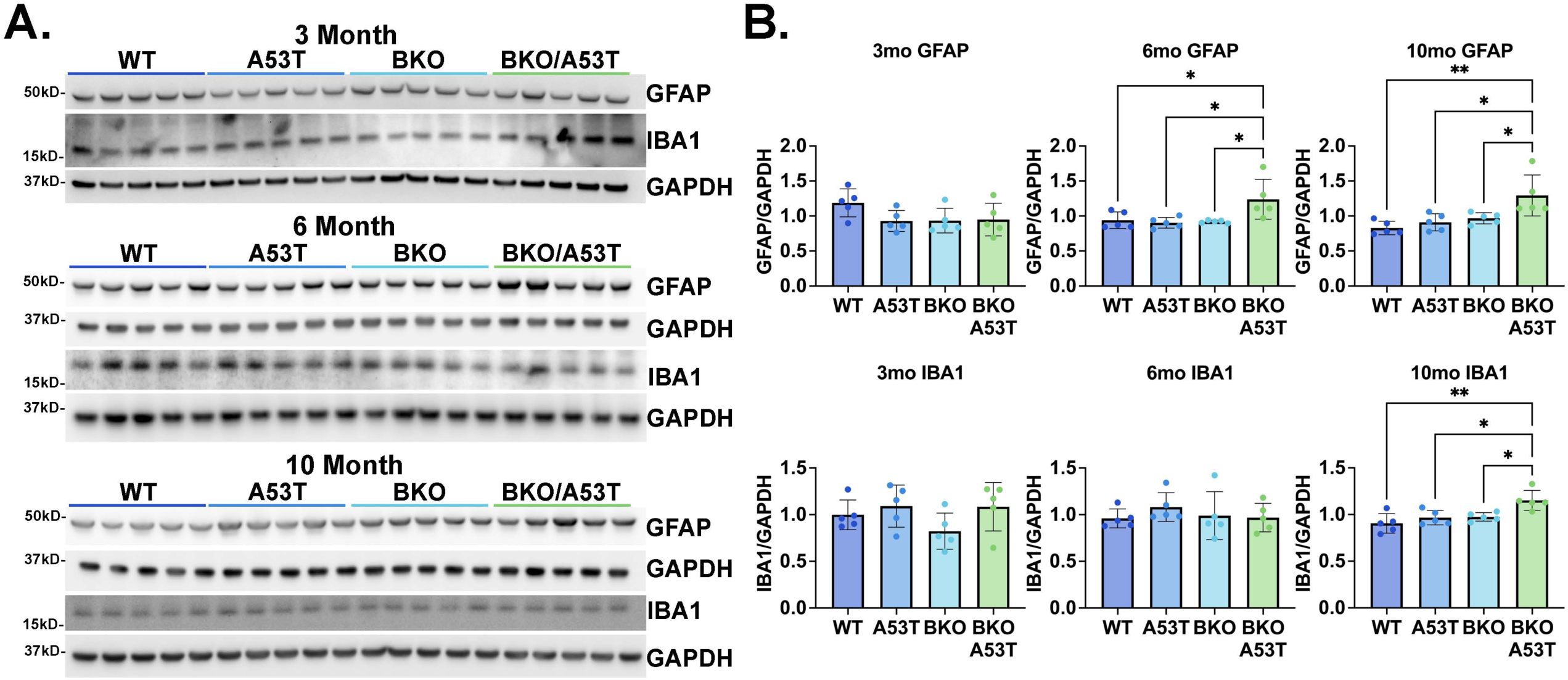
Rab27b KO/A53T^+^ mice show elevated GFAP and Iba1 in cortical lysates. **a:** Representative Western blots for GFAP, Iba1, and GAPDH in cortical lysates from three-, six-, and 10-month-old WT, Rab27b KO, A53T^+^, and Rab27b KO/A53T^+^ mice. b: Quantification of GFAP (top) and Iba1 (bottom) levels in cortical lysates from three-, six-, and 10-month-old WT, Rab27b KO, A53T^+^, and Rab27b KO/A53T^+^ mice. One-way ANOVA followed by Tukey’s multiple comparisons post-hoc test; *p<0.05, **p<0.01. N=5 individual mice per group. All error bars denote standard deviation from the mean. Note that the six and 10 month GAPDH bands overlap between this figure and Fig. 3, as markers were probed on the same membranes.

We next examined the impact of Rab27b KO on microglial activation in the A53T^+^ mouse model, which does not typically exhibit microglial changes beyond the hindbrain^44,45^. Iba1 levels were comparable between all groups at six months of age when measured by either immunohistochemistry or Western blotting (Fig. S2b, 8a-b). At 10 months, however, Rab27b KO/A53T^+^ mice expressed mildly elevated Iba1 in cortical lysates compared to all other groups by Western blot analysis (Fig. 8a-b). We validated this finding via IHC in our terminally aged cohort and found elevated Iba1 signal in the Ctx and SNc of Rab27b KO/A53T^+^ mice compared to all other groups (Fig. 9a,c). Also in this cohort, Rab27b KO/A53T^+^ mice showed elevated Iba1 signal in the Str relative to A53T^+^ mice (Fig. 9b).

**Figure 9:**
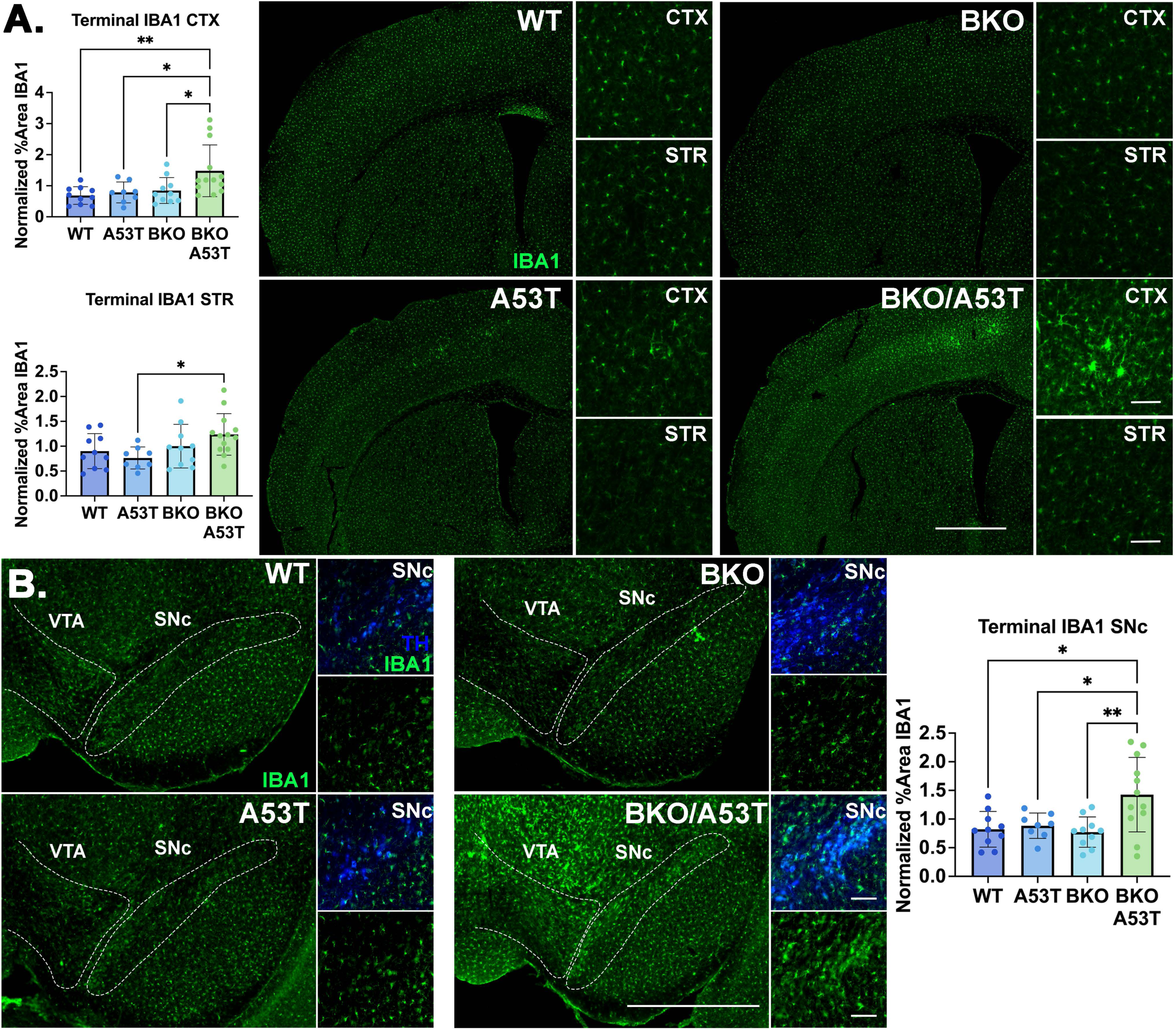
Rab27b KO/A53T^+^ mice show elevated Iba1 signal in tissue from terminally aged mice. **a:** Quantification and representative images showing Iba1 immunoreactivity in the Ctx and Str of terminally aged WT, Rab27b KO, A53T^+^, and Rab27b KO/A53T^+^ mice. Small panels are zoomed images. One-way ANOVA followed by Tukey’s multiple comparisons post-hoc test; *p<0.05, **p<0.01. N=8-13 individual mice per group. b: Quantification and representative images showing Iba1 immunoreactivity in the SNc of terminally aged WT, Rab27b KO, A53T^+^, and Rab27b KO/A53T^+^ mice. VTA and SNc were outlined using TH staining (not pictured). One-way ANOVA followed by Tukey’s multiple comparisons post-hoc test; *p<0.05, **p<0.01. N=8-13 individual mice per group. Ctx = Sensorimotor Cortex; Str = Dorsolateral Striatum; SNc = Substantia Nigra pars Compacta; VTA = Ventral Tegmental Area. Scale bars on full-sized images are 1000µm. Scale bars on enlarged images are 100µm. All error bars denote standard deviation from the mean. Note that GFAP and Iba1 were stained on the same sections (see Fig. 7).

### Rab27b KO does not induce striatal Tyrosine Hydroxylase loss in the A53T^+^ model, but does exacerbate dopaminergic loss in an AAV-αsyn overexpression model

A53T^+^ transgenic mice do not typically exhibit dopaminergic neuron loss in the SNc or dopaminergic terminal loss in the Str^40,44,46^. To test if Rab27b KO may promote dopaminergic neurodegeneration, we measured Tyrosine Hydroxylase (TH) loss in the dorsolateral Str in the terminally aged cohort. TH intensity in the dorsolateral Str was similar between all groups (Fig. S3a).

To determine if Rab27b KO could impact dopaminergic neuron loss in an alternative αsyn model, we assessed the role of Rab27b KO *in vivo* using an adeno-associated virus (AAV) αsyn overexpression model^37^. This AAV model characteristically demonstrates dopaminergic neuron loss in the SNc. Rab27b KO and WT mice were injected with AAVs expressing either WT human αsyn (rAAV-CBΑ-SYNUCLEIN-IRES-EGFP-WPRE) or GFP (rAAV-CBA-IRES-EGFP-WPRE) at two to three months of age and were sacrificed six months post injection (Fig. 10a). Injection targeting was validated via staining (Fig. 10b). We assessed TH intensity in the dorsolateral Str by immunohistochemistry. Both WT and Rab27b KO mice injected with AAV-SYN showed significant TH loss in the Str relative to WT and Rab27b KO mice injected with AAV-GFP, yet there were no differences in striatal TH intensity between WT mice injected with AAV-SYN and Rab27b mice injected with AAV-SYN (Fig. S3b). We next assessed TH-positive neuronal counts in the SNc using stereology. TH-positive neuronal counts were reduced in Rab27b KO mice injected with AAV-SYN compared to both GFP-injected groups, but WT mice injected with AAV-SYN did not display significant TH neuron loss in the SNc (Fig. 10c). These findings point to Rab27b loss accelerating dopamine neuron loss in the AAV αsyn model.

**Figure 10:**
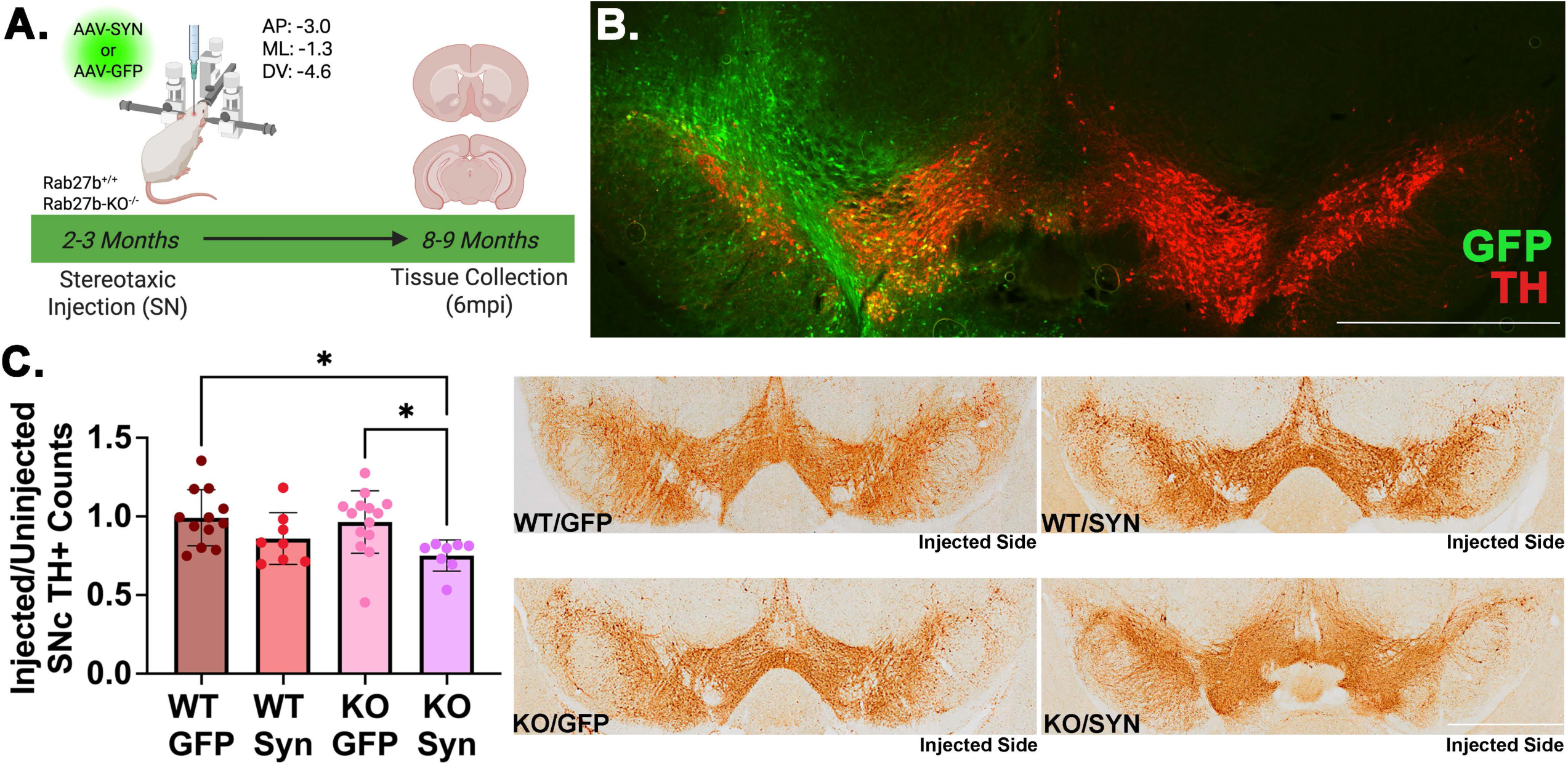
Rab27b KO accelerates TH loss in the SNc of mice injected with AAV **α**syn. a: Schematic showing experimental design. b: Representative image showing GFP expression in the SNc. c: Quantification and representative images of TH-positive neuronal counts in the SNc of WT and Rab27b KO mice injected with either AAV-GFP or AAV-SYN virus into the SNc. One-way ANOVA followed by Tukey’s multiple comparisons post-hoc test; *p<0.05. N=8-14 individual mice per group. Scale bars are 1000µm. All error bars denote standard deviation from the mean.

## DISCUSSION

Our findings show that the loss of Rab27b exacerbates neuropathology *in vivo* in two different αsyn overexpression models, highlighting Rab27b’s ability to influence pathological progression in the context of αsyn stress. We have previously shown that loss of Rab27b impairs lysosomal activity and promotes αsyn toxicity *in vitro*. Here, we show that Rab27b KO not only results in autophagic-lysosomal changes but also increases the accumulation of insoluble αsyn and neuroinflammation across multiple PD-relevant brain regions in the A53T αsyn model *in vivo*. Additionally, Rab27b KO exacerbates dopaminergic neuron loss in the AAV αsyn model.

We have previously shown that Rab27b plays a critical role in lysosomal degradative function in mouse brains. Rab27b KO mice show defects in lysosomal enzyme maturation and lysosomal degradation capabilities^35^. Similarly, here we observed that Rab27b KO disrupts autophagic-lysosomal markers in the A53T^+^ model: both the autophagic substrate marker p62 and the lysosomal marker LAMP1 were elevated in the Ctx in Rab27b KO/A53T^+^ mice compared to controls. Elevated p62 is widely regarded as a marker of generalized autophagic-lysosomal dysfunction, while LAMP1 may be elevated in a compensatory capacity. While we have previously shown that both Rab27bKO and Rab27b KO/A53T^+^ primary neurons exhibit lysosomal enzymatic defects relative to WT and A53T^+^ mice, primary neurons from Rab27b KO/A53T^+^ animals do not show exacerbated lysosomal impediments relative to Rab27b KO neurons^35^. Here we find that Rab27b KO/A53T^+^ mice show alterations in lysosomal markers compared to A53T^+^ mice and Rab27b KO mice alone *in vivo* at aged time points. This observed autophagic-lysosomal dysfunction is likely a key driver of the exacerbated neuropathology seen in Rab27b KO/A53T^+^ mice as animals age.

While A53T^+^ mice typically exhibit αsyn pathology in the hindbrain and spinal cord^36^, here we report that Rab27b KO expands and exacerbates this pathology across the Ctx, Str, and SNc, regions of relevance in PD and DLB. We measured insoluble αsyn using PK digestion across multiple sample types with slightly variable results. While both lysate and tissue PK digestion revealed elevated PK-insoluble αsyn in Rab27b KO/A53T^+^ animals relative to all other groups, only the lysate approach also showed elevated insoluble αsyn in the cortex of A53T^+^ animals alone relative to non-transgenic controls. This discrepancy for A53T^+^ mice may be driven by sensitivity differences across our assays. Assessing pS129 αsyn again revealed exacerbated pathology in Rab27b KO/A53T^+^ mice relative to A53T^+^ mice in both the Ctx and SNc, demonstrating that Rab27b KO exacerbates αsyn pathology in PD-relevant brain regions.

Interestingly, to our knowledge cortical αsyn pathology is not classically reported in this strain of A53T^+^ mice^36^. We were able to detect not only PK-resistant total αsyn, but also PK-resistant pS129-positive αsyn in the cortex of A53T^+^ animals. Although A53T^+^ animals displayed significantly less pathology than Rab27b KO/A53T^+^ animals, it is possible that newer antibodies and techniques for assessing insoluble αsyn allowed us to visualize previously unreported pathology in the A53T^+^ mouse alone. Alternatively, this difference may have arisen as the A53T^+^ line was bred and maintained over time.

In addition to increased αsyn pathology, we observed increased neuroinflammation in Rab27b KO/A53T^+^ mice. While A53T^+^ mice did not show significant changes in GFAP or Iba1 in the Ctx, Str, or SNc, Rab27b KO/A53T^+^ mice showed elevated signal for both markers in these brain regions. As the overexpression of A53T αsyn is mostly restricted to neuronal populations in this model^36^, glial activation is most likely a paracrine response to neuronal αsyn. However, KO of Rab27b is not limited to neurons but also occurs in glial populations in the Rab27b KO model. Therefore, it is not clear if this increase in neuroinflammation related to Rab27b KO is driven by potential lysosomal dysfunction in neurons and/or in astrocytes and microglia themselves. Future studies could test whether Rab27b expression in neurons vs glial cells is more important to reducing αsyn pathology and neuroinflammation in the A53T^+^ model by examination of conditional Rab27b KO in neuronal compared to glial populations.

Although the A53T^+^ mouse strain used in this study robustly overexpresses mutant αsyn throughout all brain regions, it does not typically exhibit αsyn or glial pathology outside of the hindbrain and spinal cord^36^. Here we observed significantly elevated levels of insoluble and/or pS129-positive αsyn in the Ctx, Str, and SNc, brain regions more relevant to human PD and DLB. We also saw that loss of Rab27b promoted the emergence of glial pathology in the SNc, Ctx, and Str. Given our previous work demonstrating how Rab27b is critical to lysosomal degradative function, our data here support the importance of lysosomal function to the brain’s ability to handle proteostatic stress from αsyn aggregation. We find here that damaging cells’ lysosomal capacity confers vulnerability in regions more relevant to human disease in the A53T^+^ model. This aligns with the finding that mutations in lysosomal genes and resulting lysosomal dysfunction confer elevated PD risk in human populations. Indeed, Kim *et al*. have shown that expression of mutant *GBA1* in the A53T^+^ model similarly promotes pathology in the SNc, resulting in pS129 αsyn accumulation, neuroinflammation, and dopaminergic cell loss^44^. Qualitatively, in Rab27b KO/A53T^+^ mice, both αsyn and glial pathology seem enriched in cortical Layer V, which has been identified as a selectively vulnerable region in both human brains and preformed fibril injection mouse models^47–50^. In the future, this animal model could be useful for further studies assessing both the effects of generalized lysosomal dysfunction and selectively vulnerable neuronal populations in the context of αsyn stress.

Despite increased αsyn and glial pathology in the Ctx and SNc in Rab27b KO/A53T^+^ mice, we did not observe differences in survival or wire hang capacity between A53T^+^ and Rab27b KO/A53T^+^ mice, although both αsyn transgenic groups displayed diminished wire hang capacity relative to controls. It is possible that this behavior assay was not sensitive enough to detect subtle differences in motor capacity. We also noted that both A53T^+^ and Rab27b KO/A53T^+^ mice showed reduced weights relative to non-transgenic controls across most of the aged cohort time points. A53T^+^ animals classically exhibit hyperactivity beginning around six months of age^51,52^, and this may drive the observed weight differences. Interestingly, older Rab27b KO mice alone showed diminished wire hang capacity compared to WT mice, pointing to a role for Rab27b in motor function. This finding suggests that impairment of lysosomal function due to Rab27b loss could affect motor function with age. Future work will employ a more comprehensive behavioral paradigm to examine other potential effects of Rab27b loss on motor and cognitive behavior.

We assessed TH loss in the Str of terminally aged A53T and Rab27bKO/A53T^+^ mice and did not observe changes in either model relative to controls. This is not entirely surprising, as the A53T mouse model used here does not typically exhibit dopaminergic cell loss^36^. The neuropathology we observed in the SNc may not have been enough to induce dopaminergic cell loss at the ages we examined, although Rab27b KO was sufficient to drive neuroinflammation and αsyn pathology into the SNc. In order to better assess the effects of Rab27bKO on dopaminergic cell loss, we employed an AAV-mediated αsyn overexpression model that is known to experience TH-positive cell loss in the SNc and concurrently in the Str^37^. We found that AAV-SYN injected Rab27bKO mice showed significant TH-positive neuron loss in the SNc relative to both AAV-GFP control groups, while AAV-SYN injected WT mice did not. Despite this, both WT and Rab27bKO mice exhibited reduced TH signal in the Str. In this model, unhealthy dopaminergic cells typically display TH loss in the terminals before the cell bodies themselves. Rab27b KO mice may therefore be exhibiting accelerated pathology relative to WT controls.

## CONCLUSIONS

In conclusion, knockout of Rab27b exacerbates neuropathology in two αsyn models *in vivo*. These findings highlight the importance of targeting lysosomal function as a therapeutic strategy in synucleinopathies.

## MATERIALS AND METHODS

### Animals

All animals used in this study were maintained and used in compliance with the guidelines set forth by the National Institutes of Health (NIH) and the Institutional Animal Care and Use Committee (IACUC) protocols approved by the University of Alabama at Birmingham (UAB). Both male and female animals were used.

### Rab27b Knockout Mouse Model

Rab27b KO mice were generated as previously described^35^. Briefly, Rab27a and Rab27b double knockout animals from Miguel Seabra^53^ were crossed with wild-type C57BL/6-J (000664) mice from Jackson Laboratories for several generations until the desired Rab27b^+/-^ heterozygous animals were obtained and crossed to each other to generate litters with both WT (Rab27b^+/+)^ and Rab27b KO (Rab27b^-/-^) animals for experimental use.

### Rab27b Knockout A53T Transgenic Mouse Model

Rab27b KO/A53T^+^ mice were generated as previously described^35^. Briefly, Rab27b KO mice from the line described above were crossed with hemizygous A53T^+^ (A53T^+/-^) animals from Jackson Laboratories (G2-3 strain; 006823)^36^. Two parallel final crosses were established: Rab27b WT/A53T^+^ (Rab27b^+/+^ A53T^+/-^ x Rab27b^+/+^ A53T^-/-^) and Rab27b KO/A53T^+^ (Rab27b^-/-^A53T^+/-^ x Rab27b^-/-^ A53T^-/-^). Littermates from these two separate lines were collected and pooled to generate final cohorts comprised of four final genotypes: WT (Rab27^+/+^ A53T^-/-^), A53T^+^ (Rab27b^+/+^ A53T^+/-^), Rab27b KO (Rab27b^-/-^ A53T^-/-^), and Rab27b KO/A53T^+^ (Rab27b^-/-^A53T^+/-^). See Figure 1 for schematic.

### AAV Injections

#### Viral Constructs

AAV-GFP (rAAV-CBA-IRES-EGFP-WPRE) and AAV-SYN (rAAV-CBΑ-SYNUCLEIN-IRES-EGFP-WPRE) constructs from Dr. David Standaert^37^ were packaged at the University of Iowa Viral Vector Core. Viral titers were estimated using ddPCR. Titer for AAV-GFP was 1.10×10^12^ TU/ml, and titer for AAV-SYN was 1.04×10^12^ TU/ml.

#### Stereotaxic Injections

Rab27b WT and Rab27b KO mice were unilaterally injected with either AAV-SYN to overexpress alpha-synuclein or AAV-GFP to express GFP. Surgeries were performed at 8-12 weeks of age, and intracranial stereotaxic injections were targeted to the Substantia Nigra pars Compacta (SNc; coordinates from bregma AP −3.0, ML −1.3, DV −4.6). Animals were deeply anesthetized with 2-5% isoflurane and supplemental oxygen using an isoflurane vaporizer with a non-rebreathing circuit. Injections of 2µl of each virus were performed with a microinjection pump (Stoelting 10-000-004) and syringe (Hamilton 80030) at a rate of 0.25µl per minute. Animals were sacrificed at six months post injection (mpi), and targeting accuracy was validated via immunohistochemistry (Fig. 10b).

### Tissue Collection

#### Perfusion and Sectioning

Three-month-old, six-month-old, and terminally aged animals were deeply anesthetized with isoflurane before undergoing pericardial perfusion with PBS followed by 4% paraformaldehyde (PFA). Collected brains were post-fixed in 4% PFA at 4°C before undergoing cryoprotection in 30% sucrose in Phosphate Buffered Solution (PBS) and flash-freezing in 2-methylbutane. Brains were sectioned at 30µm on a sliding microtome (Leica SM2010 R) and stored at −20°C in 50% glycerol/50% PBS as a cryoprotectant.

#### Lysate Preparation

Three-, six-, and 10-month brains were homogenized in lysis buffer (50mM Tris-HCL pH 7.4, 175mM NaCl, and 5mM EDTA) using a motorized tissue grinder and then sonicated using a probe-tip sonicator. To generate a total protein fraction, 100ul of lysate was set aside and incubated on ice with the addition of 1% Triton X-100 and 0.1% sodium dodecyl sulfate (SDS) for 30 minutes. The remaining lysate was incubated in 1% Triton X-100 for 30 minutes on ice and then centrifuged at 15,000xg for 60 minutes. The resulting supernatant was collected as the Triton X-100-soluble fraction. The pellet was resuspended in lysis buffer containing 2% SDS, sonicated, and centrifuged at 15,000g for 10 minutes to generate the SDS-soluble fraction. After protein estimation by BCA, a 20µg aliquot of protein from the SDS fraction was digested with 50µg/ml proteinase K (PK) at 37°C for 15 minutes to generate the PK-resistant fraction^54,55^.

#### Survival Analysis

Survival cohort animals were sacrificed via pericardial perfusion as described above upon the onset of severe hind-limb paralysis. Severe paralysis was determined when the hind limbs were visibly dragging and the animal was unable to reliably rise to access food and water. Investigators were blinded to animal identity during survival monitoring.

#### Wire Hang Behavioral Assay

Animals were placed on a four-paw wire hang apparatus created by the UAB machine shop. The apparatus was suspended over a rat cage with excess bedding at a height of 24 inches. The latency to fall was recorded across two trials with a one-hour rest period between trials. The maximum hold time for each animal was used. Animals were weighed before the first trial, and weights were used to calculate the maximum holding impulse [weight (g) * maximum hold time (s)], which is used to normalize for any differences in mouse weights^41^.

### Immunohistochemistry

#### Fluorescent immunohistochemistry

Sections were washed once in Tris-buffered saline (TBS) before undergoing antigen retrieval at 37°C for one hour in antigen retrieval buffer (10mM Na Citrate, 0.05% Tween-20, pH 6.0). Tissue was washed in TBS before permeabilization with 0.25% Triton X-100 for 20 minutes at room temperature. Tissue was then blocked in 5% normal goat serum (NGS) with 0.1% Triton X-100, washed once in TBS, and then incubated in primary antibody diluted in 5% NGS at 4°C overnight. Tissue was washed before being incubated in secondary antibody at 1:500 in 5% NGS for one hour. After washing, tissue was mounted, allowed to dry, and coverslipped with Prolong Diamond with DAPI (Invitrogen P36962). Stained tissue was imaged on either an Olympus VS200 slide scanning microscope at 20x magnification or a Nikon AXR confocal microscope with NSPARC detection at 20x magnification. See Table 1 for antibody details.

**Table 1.** Antibodies.

| Primary Antibody | Dilution | Source | Identifier |
| --- | --- | --- | --- |
| **αsyn** | WB: 1:1000<br>WB 1:500 (PK-digestion)<br>IHC: 1:100 | BD Biosciences | 610787 |
| **GAPDH** | WB: 1:5000 | Cell Signaling | 14C10 |
| **GFAP** | WB: 1:1000<br>IHC: 1:1000 | Cell Signaling | 3670S |
| **GFP** | IHC: 1:300 | Invitrogen | MA5-15256 |
| **Iba1** | IHC: 1:1000 | Fujifilm | 019-19741 |
| **Iba1** | WB: 1:500 | Abcam | 5076 |
| **LAMP1** | WB 1:250 | Iowa DSHB | 1D4B |
| **p62** | WB: 1:1000 | Abcam | Ab56416 |
| **pS129** | IHC 1:1000 | Abcam | Ab51253 |
| **TH** | IHC: 1:1000 | Pell-Freeze | P40101-150 |
| **TH** | IHC: 1:2000<br>IHC: 1:100 (PK-digestion) | Sigma | AB1542 |

**Table 2.** Terminally aged cohort demographics.

| Genotype | N | Age Range (Months) | Average Age ± StDev |
| --- | --- | --- | --- |
| **WT (Rab27<sup>+/+</sup> A53T<sup>-/-</sup>)** | 10 | 9.4-16.6 | 13.2 ± 2.8 |
| **A53T<sup>+</sup> (Rab27b<sup>+/+</sup> A53T<sup>+/-</sup>)** | 8 | 8.1-18.7 | 14.1 ± 3.6 |
| **Rab27b KO (Rab27b<sup>-/-</sup> A53T<sup>-/-</sup>)** | 10 | 9.8-17.3 | 14.9 ± 2.8 |
| **Rab27b KO/A53T<sup>+</sup> (Rab27b<sup>-/-</sup> A53T<sup>+/-</sup>)** | 13 | 9.2-18.0 | 14.4 ± 2.9 |

#### Proteinase K Digestion

Tissue was mounted onto slides and allowed to dry overnight. Wells were split across two slides, and a hydrophobic barrier was drawn around the sections. For each animal, one slide was incubated in 50µg/ml PK in TBS for 10 minutes at 37°C, and a control slide was incubated in TBS for 10 minutes at 37°C. After digestion, staining proceeded as described above, and slides were imaged on an Olympus VS200 slide scanning microscope at 20x magnification.

#### Stereological Analysis

For stereological analysis, the midbrain was sectioned and stored serially across five wells per animal. Sections were quenched in 0.6% hydrogen peroxide in methanol for 20 minutes before undergoing antigen retrieval as described above. Sections were blocked in 5% NGS and 0.3% Triton X-100 in TBS for one hour and incubated in primary antibody diluted in 1% NGS and 0.3% Triton X-100 overnight. Tissue was washed before undergoing a four-hour incubation at 4°C in secondary antibody diluted 1:500 in 2.5% NGS and 0.3% Triton X-100 in TBS. After washing, sections were incubated in ABC-HRP solution (VectorLabs PK-4000) for 30 minutes and developed in ImmPACT DAB Substrate (VectorLabs SK-4105). Sections were mounted on slides and dried overnight before dehydration with ethanol and Histo-Clear followed by coverslipping with Permount (Fisher Chemical SP15-100). See Table 1for antibody details.

Stereological analysis was performed using StereoInvestigator’s optical fractionator (MBF Biosciences, version 2023.2.3) as previously described^56,57^. Briefly, SNc regions of every fifth section were outlined, and 50µm x 50µm counting frames were randomly assigned by StereoInvestigator. Within each frame, tyrosine hydroxylase (TH)-positive nuclei were counted using an Olympus BX51 Microscope with a 60x oil immersion objective. Nuclei were counted within three-dimensional optical dissectors set to 20µm. Section thickness was measured at the beginning of each animal, and the total number of neurons was calculated using the following equation: Total Neurons = Neurons Counted x [1 / Section Sampling Fraction] x [1 / Area Sampling Fraction] x [1 / Height Sampling Fraction]. Both injected and uninjected sides were counted, and injected side counts were normalized to the uninjected side within each animal. Investigators were blinded to experimental conditions.

#### Western Blotting

Western blotting was performed as previously described^58^. Briefly, 20µg of protein was denatured by boiling in sample loading buffer (0.25M Tris-HCl pH 6.8, 8% SDS, 200mM DTT, 30% Glycerol, bromophenol blue dye) for 10 minutes. Samples were loaded into either 15% bis-acrylamide (BioRad 1610158) gels or 4-12% gradient NuPAGE Bis-Tris Midi gels (Invitrogen WG1402BOX). Membranes were fixed in 0.4% PFA for 30 minutes before blocking and antibody incubations. See Table 1 for antibody details. Blots were imaged using a ChemiDoc MP system (Bio-Rad 12003154) and analyzed in ImageLab (Bio-Rad version 6.1.0).

## Statistical Analyses

Statistical analyses were performed using GraphPad Prism 10 (version 10.4.2). See Table S1 for details and summary statistics.

## DATA AVAILABILITY

The datasets generated and analyzed for this study are available from the corresponding author upon request.

## AUTHOR CONTRIBUTIONS

KS designed and performed experiments, analyzed data, and wrote the manuscript. MAG, LM, RS, WJS, CC, and RE performed assays, analyzed data, and reviewed the manuscript. TY designed assays and edited and finalized the manuscript.

## ACKNOWLEDGEMENTS

This study was supported by NIH [RF1NS115767 (TAY); P50 NS108675 (TAY); T32 5T32GM135028 (KS); T32 5T32NS095775 (KS); F31 1F31NS135928-01A1 (KS)] and the American Parkinson Disease Association. Behavioral studies were facilitated by the UAB Behavioral Core Facility. We would additionally like to thank the UAB ADRC Neuropathology Core for the use of the Olympus VS200 microscope (P30AG086401). Graphics were made using BioRender (Scholz, K. (2026) https://BioRender.com/w8iuc9l; Scholz, K. (2026) https://BioRender.com/si2fwk4).

