## Supplemental Figure Legends for "Knockout of Rab27b exacerbates neuropathology in alpha-synuclein mouse models"

**Figure S1: Rab27b KO does not alter levels of PK-insoluble αsyn in three-month-old mice.**

a-b: Representative images (a) and quantification (b) of αsyn immunoreactivity in the Ctx and Str from brain tissue with and without PK digestion from three-month-old WT, A53T^+^, Rab27b KO, and Rab27b KO/A53T^+^ mice. One-way ANOVA followed by Tukey’s Multiple Comparisons post-hoc test; **p<0.01, ***p<0.001. N=4-5 individual mice per group.

Scale bars are 1000µm.

**Figure S2: Six-month-old Rab27b KO/A53T^+^ mice show elevated GFAP but not Iba1.**

a: Representative images and quantification of GFAP immunoreactivity in the Ctx and Str of six-month-old WT, Rab27b KO, A53T^+^, and Rab27b KO/A53T^+^ mice. Small panels are zoomed images. One-way ANOVA followed by Tukey’s Multiple Comparisons post-hoc test; *p<0.05, **p<0.01. N=4-5 individual mice per group.

b: Representative images and quantification of Iba1immunoreactivity in the Ctx and Str of six-month-old WT, Rab27b KO, A53T^+^, and Rab27b KO/A53T^+^ mice. Small panels are zoomed images. One-way ANOVA. N=4-5 individual mice per group.

Ctx = Sensorimotor Cortex; Str = Dorsolateral Striatum. Note that GFAP and Iba1 were stained on the same sections. Scale bars on full-sized images are 1000µm. Small scale bars are 100µm.

**Figure S3: Striatal TH loss was not changed by Rab27b KO in any model.**

a: Representative images and quantification of TH immunoreactivity in the Str of the terminally aged WT, Rab27b KO, A53T^+^, and Rab27b KO/A53T^+^ mice. One-way ANOVA. N=7-13 individual mice per group.

b: Representative images and quantification of TH immunoreactivity in the Str of WT and Rab27b KO mice injected with either AAV-GFP or AAV-SYN virus into the SNc. One-way ANOVA followed by Tukey’s Multiple Comparisons post-hoc test; *p<0.05, **p<0.01. N=8=14 individual mice per group.
