## Supplementary figures and images for "Knockout of Rab27b exacerbates neuropathology in alpha-synuclein mouse models"

### Supplemenal Figure 1

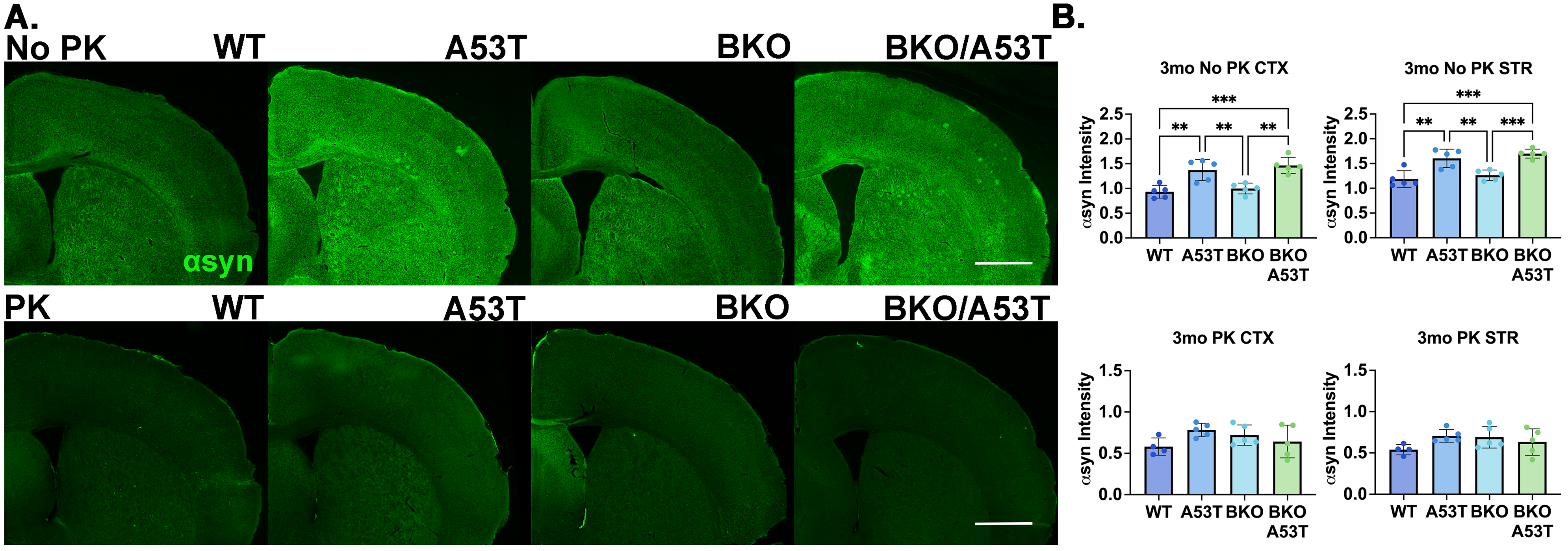

### Supplemental Figure 2

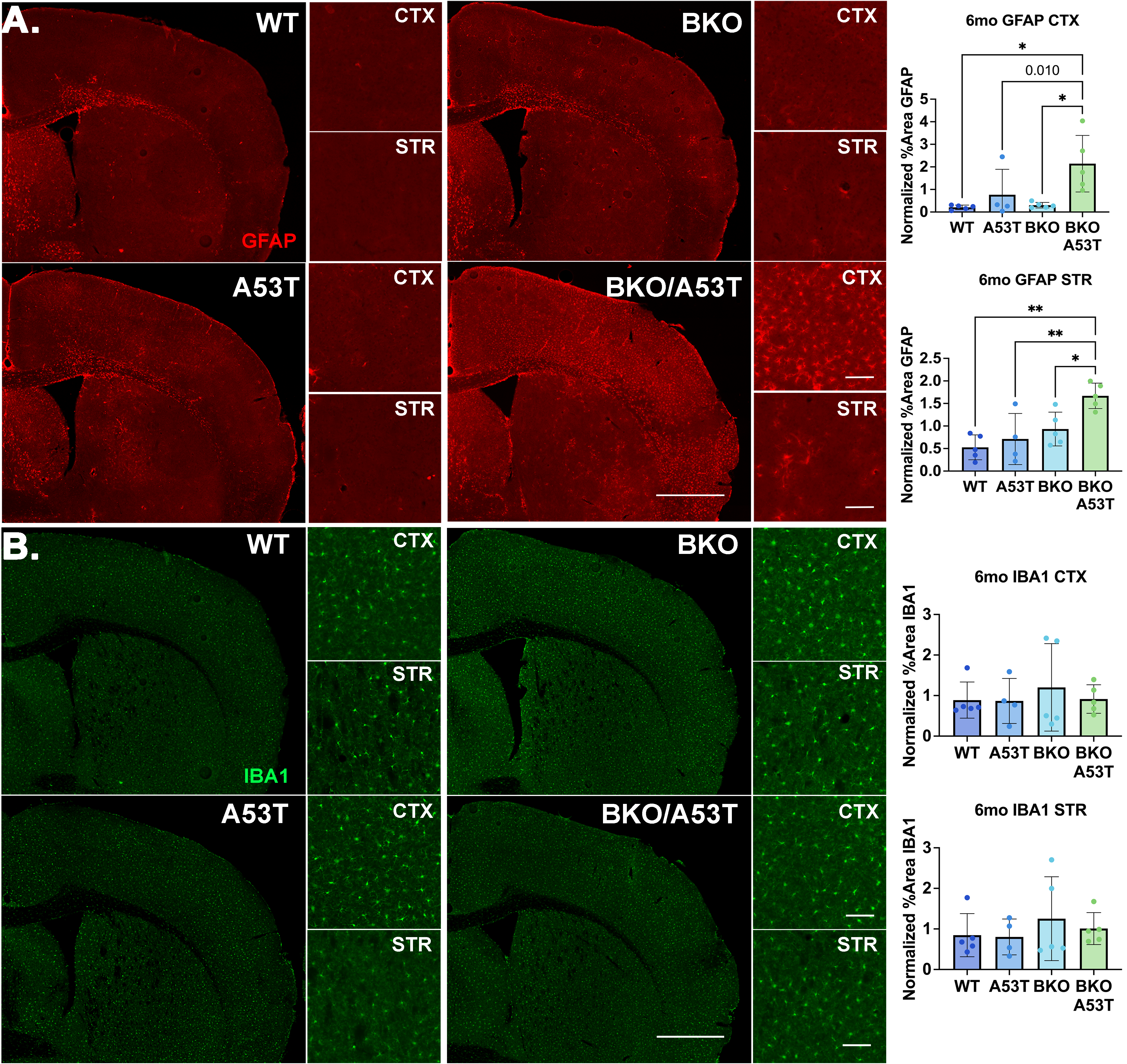

### Supplemental Figure 3

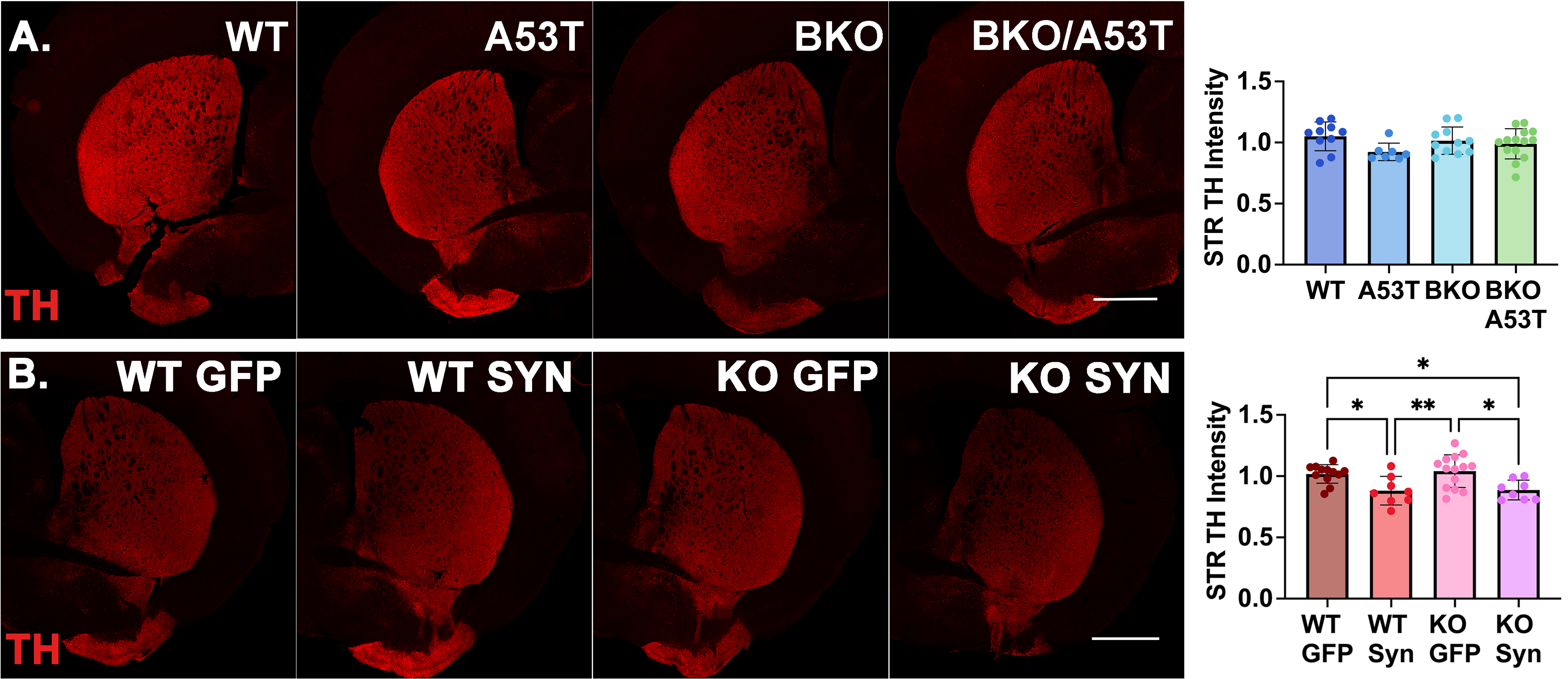
