## Supplemental Table 1 for "Knockout of Rab27b exacerbates neuropathology in alpha-synuclein mouse models"

| Fig | Expt | Sample size | Statistical analysis | Post-Hoc p-values |
| --- | --- | --- | --- | --- |
| 2A | 3-4mo Wire Hang | WT n=13 A53T n=7 BKO n=11 BKO A53T n=9 | **2-Way ANOVA** Interaction F (3, 36) = 0.7920 p=0.5064;  Genotype F (1, 36) = 0.04685 p=0.8299;  Age F (3, 36) = 2.018 p=0.1288; | **Fisher's LSD** Genotype  3mo WT vs. A53T 0.1779     3mo WT vs. BKO 0.3366     3mo WT vs. BKO A53T 0.1697     3mo A53T vs. BKO 0.619     3mo A53T vs. BKO A53T 0.9427     3mo BKO vs. BKO A53T 0.6495     4mo  WT vs. A53T 0.0291     4mo WT vs. BKO 0.3452    4mo WT vs. BKO A53T 0.0197     4mo A53T vs. BKO 0.1802     4mo A53T vs. BKO A53T 0.9853     4mo BKO vs. BKO A53T 0.1556 Age    WT  3mo vs. 4mo 0.4573     A53T 3mo vs. 4mo 0.4845     BKO 3mo vs. 4mo 0.477     BKO A53T 3mo vs. 4mo 0.3762 |
| 2B | 4-3mo Weight | WT n=13  A53T n=7  BKO n=11  BKO A53T n=9 | **2-Way ANOVA**  Interaction F (3, 23) = 3.052 p=0.0489  Genotype F (3, 23) = 0.7080 p=0.5571  Age F (1, 23) = 67.02 p<0.0001 | **Fisher's LSD**  Genotype  3mo WT vs. A53T 0.373  3mo WT vs. BKO 0.8857  3mo WT vs. BKO A53T 0.5802  3mo A53T vs. BKO 0.3068  3mo A53T vs. BKO A53T 0.7823  3mo BKO vs. BKO A53T 0.4975  4mo WT vs. A53T 0.185  4mo WT vs. BKO 0.8256  4mo WT vs. BKO A53T 0.1941  4mo A53T vs. BKO 0.26  4mo A53T vs. BKO A53T 0.968  4mo BKO vs. BKO A53T 0.2671  Age  WT 3mo vs. 4mo <0.0001  A53T 3mo vs. 4mo 0.0012  BKO 3mo vs. 4mo <0.0001  BKO A53T 3mo vs. 4mo 0.1062 |
| 2C | Survival Analysis | A53T n=7 BKO A53T n=11 | **Log-Rank (Mantel-Cox) test** Chi-square = 0.05755; df=1 p=8.104 |  |
| 2D | 7-10mo Wire Hang | WT n=13 A53T n=8-10* BKO n=10 BKO A53T n=13-15*  *animals were removed from cohort as they developed terminal paralysis and were sacrificed. | **Mixed Effects Model (REML)** Interaction F (9, 125) = 2.371 p=0.0165 Genotype F (2.299, 95.79) = 3.851 p=0.0197 Age F (3, 44) = 4.199 p=0.0107 | **Fisher's LSD** Genotype  7mo WT vs. A53T 0.013  7mo WT vs. BKO 0.0402  7mo WT vs. BKO A53T 0.0748  7mo A53T vs. BKO 0.2893  7mo A53T vs. BKO A53T 0.0104  7mo BKO vs. BKO A53T 0.3645  8mo WT vs. A53T 0.0412  8mo WT vs. BKO 0.0174  8mo WT vs. BKO A53T 0.0471  8om A53T vs. BKO 0.7774  8mo A53T vs. BKO A53T 0.6064  8mo BKO vs. BKO A53T 0.2163  9mo WT vs. A53T 0.0315  9mo WT vs. BKO 0.0516  9mo WT vs. BKO A53T 0.0497  9mo A53T vs. BKO 0.5751  9mo A53T vs. BKO A53T 0.4729  9mo BKO vs. BKO A53T 0.8993  10mo WT vs. A53T 0.2711  10mo WT vs. BKO 0.3632  10mo WT vs. BKO A53T 0.2161  10mo A53T vs. BKO 0.6725  10mo A53T vs. BKO A53T 0.8313  10mo BKO vs. BKO A53T 0.6751 Age  WT 7mo vs. 8mo 0.4209  WT 7mo vs. 9mo 0.0955  WT 7mo vs. 10mo 0.016  WT 8mo vs. 9mo 0.0698  WT 8mo vs. 10mo 0.0256  WT 9mo vs. 10mo 0.057  A53T 7mo vs. 8mo 0.1577  A53T 7mo vs. 9mo 0.3782  A53T 7mo vs. 10mo 0.6498  A53T 8mo vs. 9mo 0.2948  A53T 8mo vs. 10mo 0.4268  A53T 9mo vs. 10mo 0.9106  BKO 7mo vs. 8mo 0.3208  BKO 7mo vs. 9mo 0.8438  BKO 7mo vs. 10mo 0.8538  BKO 8mo vs. 9mo 0.4939  BKO 8mo vs. 10mo 0.5943  BKO 9mo vs. 10mo 0.9923  BKO A53T 7mo vs. 8mo 0.1708  BKO A53T 7mo vs. 9mo 0.0716  BKO A53T 7mo vs. 10mo 0.0444  BKO A53T 8mo vs. 9mo 0.1782  BKO A53T 8mo vs. 10mo 0.1956  BKO A53T 9mo vs. 10mo 0.5109 |
| 2E | 7-10mo Weight | WT n=13 A53T n=8-10* BKO n=10 BKO A53T n=13-15*  *animals were removed from cohort as they developed terminal paralysis and were sacrificed. | **Mixed Effects Model (REML)** Interaction F (9, 125) = 1.466 p=0.1676 Genotype F (3, 125) = 13.25 p<0.0001 Age F (3, 44) = 5.007 p=0.0045 | **Fisher's LSD** Genotype     7mo WT vs. A53T 0.0413     7mo WT vs. BKO 0.3638     7mo WT vs. BKO A53T 0.101     7mo A53T vs. BKO 0.0059     7mo A53T vs. BKO A53T 0.5576     7mo BKO vs. BKO A53T 0.0145     8mo WT vs. A53T 0.0135     8mo WT vs. BKO 0.4573     8mo WT vs. BKO A53T 0.0369     8mo A53T vs. BKO 0.0027     8mo A53T vs. BKO A53T 0.5319     8mo BKO vs. BKO A53T 0.0072     9mo WT vs. A53T 0.0101     9mo WT vs. BKO 0.2698     9mo WT vs. BKO A53T 0.0832     9mo A53T vs. BKO 0.0006     9mo A53T vs. BKO A53T 0.2886     9mo BKO vs. BKO A53T 0.0065     10mo WT vs. A53T 0.0061     10mo WT vs. BKO 0.1553     10mo WT vs. BKO A53T 0.1101     10mo A53T vs. BKO 0.0001     10mo A53T vs. BKO A53T 0.1712     10mo BKO vs. BKO A53T 0.0035 Age     WT 7mo vs. 8mo 0.0083     WT 7mo vs. 9mo 0.0025     WT 7mo vs. 10mo 0.0009     WT 8mo vs. 9mo 0.6913     WT 8mo vs. 10mo 0.4735     WT 9mo vs. 10mo 0.7488     A53T 7mo vs. 8mo 0.4872     A53T 7mo vs. 9mo 0.5067     A53T 7mo vs. 10mo 0.7826     A53T 8mo vs. 9mo 0.9754     A53T 8mo vs. 10mo 0.6955     A53T 9mo vs. 10mo 0.7175     BKO 7mo vs. 8mo 0.0814     BKO 7mo vs. 9mo 0.0009     BKO 7mo vs. 10mo <0.0001     BKO 8mo vs. 9mo 0.1006     BKO 8mo vs. 10mo 0.0025     BKO 9mo vs. 10mo 0.1539      BKO A53T 7mo vs. 8mo 0.2841      BKO A53T 7mo vs. 9mo 0.0051      BKO A53T 7mo vs. 10mo 0.0004      BKO A53T 8mo vs. 9mo 0.0736      BKO A53T 8mo vs. 10mo 0.0106      BKO A53T 9mo vs. 10mo 0.4099 |
| 3 | ALM Western Blots | Per time point WT n=5 A53T n=5 BKO n=5 BKO A53T n=5 | **One-Way ANOVA** 3mo P62 F (3, 16) = 2.965 P=0.0635 6mo P62 F (3, 16) = 3.240 P=0.0500 10mo P62 F (3, 16) = 5.159 P=0.0110 3mo LAMP1 F (3, 16) = 0.3866 P=0.7641 6mo LAMP1 F (3, 16) = 1.199 P=0.3420 10mo LAMP1 F (3, 16) = 20.00 P<0.0001 | **Tukey's Multiple Comparisons** 3mo P62   WT vs. A53T 0.9995   WT vs. BKO 0.5132   WT vs. BKO A53T 0.0848   A53T vs. BKO 0.5788   A53T vs. BKO A53T 0.1038   BKO vs. BKO A53T 0.6517 6mo P62   WT vs. A53T 0.9514   WT vs. BKO 0.5136   WT vs. BKO A53T 0.0455   A53T vs. BKO 0.8153   A53T vs. BKO A53T 0.1207   BKO vs. BKO A53T 0.4581 10mo P62   WT vs. A53T 0.9824   WT vs. BKO >0.9999   WT vs. BKO A53T 0.0213   A53T vs. BKO 0.9745   A53T vs. BKO A53T 0.0439   BKO vs. BKO A53T 0.01923 3mo LAMP1   WT vs. A53T 0.866   WT vs. BKO 0.9935   WT vs. BKO A53T 0.9996   A53T vs. BKO 0.7332   A53T vs. BKO A53T 0.9079   BKO vs. BKO A53T 0.9827 6mo LAMP1   WT vs. A53T 0.3948   WT vs. BKO 0.3791   WT vs. BKO A53T 0.6309   A53T vs. BKO >0.9999   A53T vs. BKO A53T 0.9753   BKO vs. BKO A53T 0.9697 10mo LAMP1   WT vs. A53T 0.0085   WT vs. BKO 0.0533   WT vs. BKO A53T <0.0001   A53T vs. BKO 0.7919   A53T vs. BKO A53T 0.0063   BKO vs. BKO A53T 0.001 |
| 4 | Asyn Western Blots | Per time point WT n=5 A53T n=5 BKO n=5 BKO A53T n=5 | **One-Way ANOVA** 3mo Triton F (3, 16) = 154.7" p<0.0001 3mo SDS F (3, 16) = 47.83" p<0.0001 6mo Triton F (3, 16) = 58.27" p<0.0001 6mo SDS F (3, 16) = 140.2" p<0.0001 6mo PK F (3, 16) = 22.19" p<0.0001 10mo Triton F (3, 16) = 123.6" p<0.0001 10mo SDS F (3, 16) = 35.69" p<0.0001 10mo PK F (3, 16) = 16.33" p<0.0001 | **Tukey's Multiple Comparisons** 3mo Triton   WT vs. A53T <0.0001   WT vs. BKO 0.249   WT vs. BKO A53T <0.0001   A53T vs. BKO <0.0001   A53T vs. BKO A53T 0.6433   BKO vs. BKO A53T <0.0001 3mo SDS   WT vs. A53T <0.0001   WT vs. BKO 0.9999   WT vs. BKO A53T <0.0001   A53T vs. BKO <0.0001   A53T vs. BKO A53T >0.9999   BKO vs. BKO A53T <0.0001 6mo Triton   WT vs. A53T <0.0001   WT vs. BKO 0.9986   WT vs. BKO A53T <0.0001   A53T vs. BKO <0.0001   A53T vs. BKO A53T 0.9675   BKO vs. BKO A53T <0.0001 6mo SDS   WT vs. A53T <0.0001   WT vs. BKO 0.7658   WT vs. BKO A53T <0.0001   A53T vs. BKO <0.0001   A53T vs. BKO A53T 0.9931   BKO vs. BKO A53T <0.0001 6mo PK   WT vs. A53T 0.0439   WT vs. BKO 0.9713   WT vs. BKO A53T <0.0001   A53T vs. BKO 0.0995   A53T vs. BKO A53T 0.0026   BKO vs. BKO A53T <0.0001 10mo Triton   WT vs. A53T <0.0001   WT vs. BKO 0.9362   WT vs. BKO A53T <0.0001   A53T vs. BKO <0.0001   A53T vs. BKO A53T 0.8048   BKO vs. BKO A53T <0.0001 10mo SDS   WT vs. A53T <0.0001   WT vs. BKO 0.9985   WT vs. BKO A53T <0.0001   A53T vs. BKO <0.0001   A53T vs. BKO A53T 0.3885   BKO vs. BKO A53T <0.0001 10mo PK   WT vs. A53T 0.0342   WT vs. BKO 0.0696   WT vs. BKO A53T <0.0001   A53T vs. BKO 0.9819   A53T vs. BKO A53T 0.0065   BKO vs. BKO A53T 0.0031 |
| S1 | 3mo PK Digestion | No PK WT n=5 A53T n=5 BKO n=5 BKO A53T n=5 PK WT n=4 (1 excluded for technical staining issue) A53T n=5 BKO n=5 BKO A53T n=5 | **One-Way ANOVA** No PK Ctx F (3, 16) = 13.95 p<0.0001 PK Ctx F (3, 15) = 1.915 p=0.1705 No PK Str F (3, 16) = 15.61 p<0.0001 PK Str F (3, 15) = 1.779 p=0.1943 | **Tukey's Multiple Comparisons** No PK Ctx   WT vs. A53T 0.0025   WT vs. BKO 0.9143   WT vs. BKO A53T 0.0004   A53T vs. BKO 0.0092   A53T vs. BKO A53T 0.7871   BKO vs. BKO A53T 0.0014 PK Ctx   WT vs. A53T 0.1646   WT vs. BKO 0.448   WT vs. BKO A53T 0.9053   A53T vs. BKO 0.8838   A53T vs. BKO A53T 0.3915   BKO vs. BKO A53T 0.8037 No PK Str   WT vs. A53T 0.0014   WT vs. BKO 0.8243   WT vs. BKO A53T 0.0002   A53T vs. BKO 0.008   A53T vs. BKO A53T 0.72   BKO vs. BKO A53T 0.0009 PK Str   WT vs. A53T 0.197   WT vs. BKO 0.2646   WT vs. BKO A53T 0.6555   A53T vs. BKO 0.9969   A53T vs. BKO A53T 0.7563   BKO vs. BKO A53T 0.8581 |
| 5A-B | 6mo PK Digestion | No PK WT n=5 A53T n=4 BKO n=5 BKO A53T n=5 PK WT n=4 (1 excluded for technical staining issue) A53T n=4 BKO n=5 BKO A53T n=5 | **One-Way ANOVA** No PK Ctx F (3, 15) = 18.39 p<0.0001 PK Ctx F (3, 14) = 7.709 p=0.0028 No PK Str F (3, 15) = 15.53 p<0.0001 PK Str F (3, 14) = 12.15 p=0.0003 | **Tukey's Multiple Comparisons** No PK Ctx   WT vs. A53T 0.017   WT vs. BKO 0.9914   WT vs. BKO A53T <0.0001   A53T vs. BKO 0.0288   A53T vs. BKO A53T 0.0824   BKO vs. BKO A53T 0.0001 PK Ctx   WT vs. A53T 0.9627   WT vs. BKO 0.7587   WT vs. BKO A53T 0.0093   A53T vs. BKO 0.4678   A53T vs. BKO A53T 0.0036   BKO vs. BKO A53T 0.0435 No PK Str   WT vs. A53T 0.0158   WT vs. BKO 0.9993   WT vs. BKO A53T 0.0002   A53T vs. BKO 0.0198   A53T vs. BKO A53T 0.2631   BKO vs. BKO A53T 0.0003 PK Str   WT vs. A53T 0.4802   WT vs. BKO 0.7014   WT vs. BKO A53T 0.0044   A53T vs. BKO 0.0811   A53T vs. BKO A53T 0.0003   BKO vs. BKO A53T 0.024 |
| 5C-D | Terminal PK Digestion (Ctx+Str) | No PK WT n=10 A53T n=8 BKO n=10 BKO A53T n=13 PK WT n=10  A53T n=7 (1 excluded for technical staining issue) BKO n=8 (2 excluded for technical staining issue) BKO A53T n=12 (2 excluded for technical staining issue) | **One-Way ANOVA** No PK Ctx F (3, 37) = 50.02 p<0.0001 PK Ctx F (3, 33) = 7.959 p=0.0004 No PK Str F (3, 37) = 74.51 p<0.0001 PK Str F (3, 33) = 7.870 p=0.0004 | **Tukey's Multiple Comparisons** No PK Ctx   WT vs. A53T <0.0001   WT vs. BKO 0.95   WT vs. BKO A53T <0.0001   A53T vs. BKO <0.0001   A53T vs. BKO A53T 0.7295   BKO vs. BKO A53T <0.0001 PK Ctx   WT vs. A53T 0.5412   WT vs. BKO 0.9913   WT vs. BKO A53T 0.0006   A53T vs. BKO 0.7418   A53T vs. BKO A53T 0.0706   BKO vs. BKO A53T 0.0031 No PK Str   WT vs. A53T <0.0001   WT vs. BKO 0.9597   WT vs. BKO A53T <0.0001   A53T vs. BKO <0.0001   A53T vs. BKO A53T 0.3443   BKO vs. BKO A53T <0.0001 PK Str   WT vs. A53T 0.391   WT vs. BKO 0.9243   WT vs. BKO A53T 0.0004   A53T vs. BKO 0.7749   A53T vs. BKO A53T 0.099   BKO vs. BKO A53T 0.0058 |
| 5E-F | Terminal PK Digestion (SNc) | No PK WT n=10 A53T n=8 BKO n=10 BKO A53T n=13 PK WT n=9 (1 excluded for technical staining issue) A53T n=7 (1 excluded for technical staining issue) BKO n=8 (2 excluded for technical staining issue) BKO A53T n=12 (2 excluded for technical staining issue) | **One-Way ANOVA** No PK SNc F (3, 37) = 40.97 p<0.0001 PK SNc F (3, 32) = 4.197 p=0.0130 | **Tukey's Multiple Comparisons** No PK SNc   WT vs. A53T <0.0001   WT vs. BKO 0.9888   WT vs. BKO A53T <0.0001   A53T vs. BKO <0.0001   A53T vs. BKO A53T 0.275   BKO vs. BKO A53T <0.0001 PK SNc   WT vs. A53T 0.999   WT vs. BKO >0.9999   WT vs. BKO A53T 0.0333   A53T vs. BKO 0.9989   A53T vs. BKO A53T 0.0747   BKO vs. BKO A53T 0.0409 |
| 6 | pS129 Area | A53T n=8 BKO A53T n=13 | **Unpaired t-test** Ctx t=2.157, df=19 p=0.0440 SNc t=2.210, df=19 p=0.0396 |  |
| 7 | Glia Western Blots | Per time point WT n=5 A53T n=5 BKO n=5 BKO A53T n=5 | **One-Way ANOVA** 3mo GFAP "F (3, 16) = 2.140" p=0.1352 6mo GFAP "F (3, 16) = 5.022" p=0.0122 10mo GFAP "F (3, 16) = 7.164" p=0.0029 3mo IBA1 "F (3, 16) = 1.731" p=0.2008 6mo IBA1 "F (3, 16) = 0.4979" p=0.6889 10mo IBA1 "F (3, 16) = 7.402" p=0.0025 | **Tukey's Multiple Comparisons** 3mo GFAP   WT vs. A53T 0.1842   WT vs. BKO 0.1981   WT vs. BKO A53T 0.2412   A53T vs. BKO >0.9999   A53T vs. BKO A53T 0.9982   BKO vs. BKO A53T 0.9993 6mo GFAP   WT vs. A53T 0.9826   WT vs. BKO 0.9968   WT vs. BKO A53T 0.041   A53T vs. BKO 0.9985   A53T vs. BKO A53T 0.0199   BKO vs. BKO A53T 0.0273 10mo GFAP   WT vs. A53T 0.8715   WT vs. BKO 0.5811   WT vs. BKO A53T 0.0027   A53T vs. BKO 0.9499   A53T vs. BKO A53T 0.0126   BKO vs. BKO A53T 0.0363 3mo IBA1   WT vs. A53T 0.8988   WT vs. BKO 0.5742   WT vs. BKO A53T 0.9144   A53T vs. BKO 0.2307   A53T vs. BKO A53T >0.9999   BKO vs. BKO A53T 0.2466 6mo IBA1   WT vs. A53T 0.7024   WT vs. BKO 0.9935   WT vs. BKO A53T 0.9998   A53T vs. BKO 0.8415   A53T vs. BKO A53T 0.7468   BKO vs. BKO A53T 0.9977 10mo IBA1   WT vs. A53T 0.6896   WT vs. BKO 0.6069   WT vs. BKO A53T 0.002   A53T vs. BKO 0.999   A53T vs. BKO A53T 0.0188   BKO vs. BKO A53T 0.0248 |
| S2 | 6mo GFAP IHC | WT n=5 A53T n=4 BKO n=5 BKO A53T n=5 | **One-Way ANOVA** Ctx F (3, 15) = 5.859 p=0.0074 Str F (3, 15) = 8.607 p=0.0015 | **Tukey's Multiple Comparisons** Ctx   WT vs. A53T 0.7436   WT vs. BKO 0.9978   WT vs. BKO A53T 0.0101   A53T vs. BKO 0.8322   A53T vs. BKO A53T 0.1034   BKO vs. BKO A53T 0.0143 Str   WT vs. A53T 0.8836   WT vs. BKO 0.3601   WT vs. BKO A53T 0.0013   A53T vs. BKO 0.821   A53T vs. BKO A53T 0.009   BKO vs. BKO A53T 0.0345 |
| 8 | Terminal GFAP IHC | Ctx WT n=10 A53T n=8 BKO n=9 (1 excluded for technical staining issue) BKO A53T n=13  Str WT n=10 A53T n=8  BKO n=9 (1 excluded for technical staining issue) BKO A53T n=12 (1 excluded for technical staining issue)  SNc WT n=10 A53T n=8 BKO n=10 BKO A53T n=12 (1 excluded for technical staining issue) | **One-Way ANOVA** Ctx F (3, 36) = 15.40 p<0.0001 Str F (3, 35) = 5.788 p=0.0025 SNc F (3, 36) = 9.413 p<0.0001 | **Tukey's Multiple Comparisons** Ctx   WT vs. A53T 0.3533   WT vs. BKO 0.9975   WT vs. BKO A53T <0.0001   A53T vs. BKO 0.4763   A53T vs. BKO A53T 0.0033   BKO vs. BKO A53T <0.0001 Str   WT vs. A53T 0.8478   WT vs. BKO 0.927   WT vs. BKO A53T 0.0031   A53T vs. BKO 0.9965   A53T vs. BKO A53T 0.0484   BKO vs. BKO A53T 0.022 SNc   WT vs. A53T 0.2164   WT vs. BKO 0.5753   WT vs. BKO A53T <0.0001   A53T vs. BKO 0.8724   A53T vs. BKO A53T 0.049   BKO vs. BKO A53T 0.0036 |
| 9 | Terminal IBA1 IHC | Ctx WT n=10 A53T n=8 BKO n=10 BKO A53T n=13  Str WT n=10 A53T n=8  BKO n=10 BKO A53T n=13  SNc WT n=10 A53T n=8 BKO n=10 BKO A53T n=12 (1 excluded for technical staining issue) | **One-Way ANOVA** Ctx F (3, 37) = 4.991 p=0.0052 Str F (3, 37) = 2.977 p=0.0439 SNc F (3, 36) = 5.760 p=0.0025 | **Tukey's Multiple Comparisons** Ctx   WT vs. A53T 0.98   WT vs. BKO 0.9132   WT vs. BKO A53T 0.0081   A53T vs. BKO 0.9956   A53T vs. BKO A53T 0.0397   BKO vs. BKO A53T 0.0467 Str   WT vs. A53T 0.868   WT vs. BKO 0.9351   WT vs. BKO A53T 0.1677   A53T vs. BKO 0.5544   A53T vs. BKO A53T 0.0396   BKO vs. BKO A53T 0.4546 SNc   WT vs. A53T 0.9896   WT vs. BKO 0.9936   WT vs. BKO A53T 0.0107   A53T vs. BKO 0.9445   A53T vs. BKO A53T 0.0396   BKO vs. BKO A53T 0.0051 |
| S3 | STR TH | A53T Cohort WT n=10 A53T n=7 (1 excluded for technical staining issue) BKO n=10 BKO A53T n=13  AAV Cohort WT GFP n=12 WT SYN n=8 KO GFP n=14 KO SYN n=8 | **One-Way ANOVA** A53T F (3, 36) = 2.077 p=0.1205 AAV F (3, 38) = 6.331 p=0.0014 | **Tukey's Multiple Comparisons** A53T   WT vs. A53T 0.0821   WT vs. BKO 0.8208   WT vs. BKO A53T 0.7983   A53T vs. BKO 0.3481   A53T vs. BKO A53T 0.3005   BKO vs. BKO A53T >0.9999 AAV   WT GFP vs. WT Syn 0.0384   WT GFP vs. KO GFP 0.9439   WT GFP vs. KO Syn 0.0484   WT Syn vs. KO GFP 0.0089   WT Syn vs. KO Syn 0.9997   KO GFP vs. KO Syn 0.0116 |
| 10 | SN TH | WT GFP n=12 WT SYN n=8 KO GFP n=14 KO SYN n=8 | **One-Way ANOVA** F (3, 38) = 3.875 p=0.0164 | **Tukey's Multiple Comparisons**   WT GFP vs. WT Syn 0.3491   WT GFP vs. KO GFP 0.9796   WT GFP vs. KO Syn 0.0202   WT Syn vs. KO GFP 0.5179   WT Syn vs. KO Syn 0.5924   KO GFP vs. KO Syn 0.0378 |
